# Drivers of inter-population variation in the gut microbiomes of sister species of *Phanaeus* dung beetles

**DOI:** 10.1101/2021.02.19.431932

**Authors:** Claire C. Winfrey, Kimberly S. Sheldon

## Abstract

The microbiome plays key roles in host physiology and ecology, but how the microbiome varies among populations of hosts is not well understood. However, different abiotic and biotic selection pressures across a species’ range likely lead to variation in the microbiome. In addition, symbiotic microbiota may differ more between closely-related species in sympatry than in allopatry if selection favors the reduction of interspecific competition. We investigated variation in the maternally-transmitted, beneficial gut microbiomes of *Phanaeus vindex* and *P. difformis*, sister species of dung beetle that compete for the same resources in sympatry and occur across diverse climatic conditions. We sampled and sequenced bacterial/archaeal 16S rDNA from guts of *P. difformis* and *P. vindex* collected across 17 sympatric and allopatric sites. Gut microbial communities were best predicted by temperature and precipitation, cattle present at sites, and spatial relationships among sites. Contrary to our hypotheses, we did not find that the gut microbial communities of *P. vindex* and *P. difformis* differed more in sympatry than in allopatry, nor that *P. vindex*, the more broadly distributed of the two species, exhibited greater microbiome turnover among populations. However, the gut microbiome of *P. vindex* shifted more between sympatric and allopatric populations than did that of *P. difformis*, suggesting character displacement. While more research is needed, it is possible that differences in gut microbial communities allow *P. vindex* and *P. difformis* to partition their niches in sympatry. Our work argues for further exploration of the gut microbiome’s potential role in niche partitioning and local adaptation.

## 1. INTRODUCTION

The unicellular organisms found in and on plants and animals, termed the microbiome, are a crucial part of the functioning of many host species. The microbiome plays diverse roles in digestion, metabolism, and immune function (Engel & Moran, 2013; Kohl & Carey, 2016), determines the thermal limits of species (Dunbar, Wilson, Ferguson, & Moran, 2007; Kikuchi et al., 2016), and may prevent hybridization (Brucker & Bordenstein, 2013). Despite the link between a host’s microbiome and its function, little is known about variation in the microbiome within and among populations of non-human hosts. However, individuals of the same species live in heterogeneous environments, and differential biotic and abiotic pressures may lead to variation in the microbiome across a species’ geographic range. Studies on species’ gut microbiomes from different populations have revealed that the gut microbiomes of lab-reared animals often differ from their wild counterparts (reviewed in Carrier & Reitzel, 2017), and importantly, that the gut microbiome of many species does vary among natural populations of the host (e.g. Coon, Brown, & Strand, 2016; Parker, Newton, & Moczek, 2020; Tiede, Scherber, Mutschler, McMahon, & Gratton, 2017). However, most previous research efforts have focused on only a few populations (but see Fietz et al., 2018; Hosokawa, et al., 2016; Wang, Kapun, Waidele, Kuenzel, Bergland, & Staubach, 2020), which are unlikely to be representative of the breadth of environmental conditions across the host’s range that may be important in structuring gut microbial communities. Thus, we still lack an understanding of the patterns and processes leading to variation in the gut microbiome of species.

One factor that could lead to variation in the gut microbiome is competition among closely-related taxa. Where ecologically-similar species occur in sympatry, natural selection should favor the species’ use of different resources or microhabitats to diffuse competition, which facilitates coexistence (Brown & Wilson, 1956; Hutchinson, 1959; MacArthur, 1958; Schoener, 1974). In contrast, ecologically-similar species occurring in allopatry are predicted to have greater niche overlap than in sympatry because of reduced competition (Brown & Wilson, 1956; Schoener, 1974). If niche partitioning underlies variation in aspects of the niche that are known to affect gut microbial communities, such as temperature or diet, then species with similar niches may have more similar gut microbiota in allopatry where their niches are more similar than in sympatry where their niches show greater divergence. To our knowledge, no studies have investigated how the presence or absence of ecologically-similar species may shape the gut microbiome across species ranges.

Alternatively, different abiotic conditions across the range of the host may drive variation in the gut microbiome by acting directly on microbes or by influencing host phenotype that in turn exerts pressure on microbes. For example, aspects of host diet that may vary across geography, such as salinity (Hallali et al., 2018; Wilck et al., 2017), fiber content (Friedman et al., 2017), and prey diversity (Tiede et al., 2017), affect gut community composition by favoring some microbes over others. Temperature variation across the range of the host may also select for distinct gut microbiomes across host populations. First, varying temperatures can alter competitive dynamics among microbes in the gut, changing the relative abundances of microbial community members (Palmer-Young, Raffel, & McFrederick, 2018). Second, gut-specialized microorganisms often have narrower thermal tolerances than their hosts (Corbin, Heyworth, Ferrari, & Hurst, 2017; Kikuchi et al., 2016; Zhang et al., 2019) or may confer different selective advantages to the host depending on the temperature (Russell & Moran, 2006). Thus, the gut microbial community may turnover across a hosts’ range based on local climatic conditions.

Variation in gut microbial communities may also be due to deterministic and stochastic processes that positively correlate with distance. Whether horizontally or vertically transmitted from one generation of hosts to the next, all gut microbes were acquired from the environment at some point in their evolutionary history (Bright & Bulgheresi, 2010), and free-living (Dickey et al., 2020; Lacap, Lau, & Pointing, 2011; Martiny et al., 2006) and host-associated (Dickey et al., 2020) microbes are subject to stochastic distance-decay processes such as dispersal limitation. By influencing the microbes that hosts consume in their diets, dispersal limitation can drive differences in the gut microbiome among populations of hosts, as observed in several species of mammal (Moeller et al., 2013; Moeller et al., 2017). Furthermore, the gut microbiome of animals is under a degree of host genetic control (e.g. Benson et al., 2010); thus, as host populations become more genetically distinct with increasing distance, so too might their gut microbiota (Fietz et al., 2018). Finally, ecological drift may also lead to increasing gut microbial dissimilarity among host populations over time (Bordenstein & Theis, 2015; Shafquat, Joice, Simmons, & Huttenhower, 2014).

We used two closely-related species of dung beetles in the genus *Phanaeus* to investigate the patterns and processes structuring the composition of the gut microbiome. *Phanaeus* develop from egg to adult completely within a ball of dung, or brood ball, that females form below dung pats (Price & May, 2009). Female dung beetles directly transmit their gut microbiome to their offspring via a bacterial pedestal within the brood ball (Estes et al., 2013). Once established in the larval gut, gut bacteria buffer the organisms against heat and desiccation stresses (Schwab, Riggs, Newton, & Moczek, 2016) and are predicted to fix nitrogen and facilitate the breakdown of cellulose (Shukla, Sanders, Byrne, & Pierce, 2016). This accelerates larval growth and development rates (Schwab et al., 2016), which are fitness proxies in insects (Kingsolver & Huey, 2008). Because the gut microbiome is vertically-transmitted, disruptions to the larval gut microbiome can cause the next generation of offspring to reach a smaller adult size (Parker, Dury, & Moczek, 2018). Therefore, the gut microbiome should be under strong selection.

*Phanaeus* vindex (MacLeay) and *P. difformis* (LeConte) are sister species of dung beetle that are morphologically near-identical and share a diet of large mammal dung (Blume & Aga, 1978; Edmonds, 1994). *P. vindex* ranges throughout most of the United States east of the Rocky Mountains, and additionally some populations can be found in desert parts of New Mexico and Arizona (Edmonds, 1994; Sheldon & Tewksbury, 2014). It is noticeably absent in southern Texas (Blume & Aga, 1978; Edmonds 1994). Although previous work suggested *P. vindex* was absent from central Texas (Blume & Aga, 1978; Edmonds 1994), in the present study we found *P. vindex* at two locations in central Texas. In contrast to *P. vindex*, *P. difformis* has a comparatively small range; it is found in Texas and Oklahoma, southern Kansas, and parts of adjacent US and Mexican states (Blume & Aga, 1978; Edmonds, 1994). Prior studies suggest that *P. difformis* prefers sandy soils, whereas *P. vindex* is found on various soil types, including clay (Dickey, 2006). Since competition for brood ball burial space is thought to be a primary force shaping dung beetle community structure and breeding and feeding behavior (Hanski & Cambefort, 1991; Price & May, 2009; Simmons & Risdill-Smith, 2011), different edaphic preferences may play a part in facilitating sympatric co-existence (Blume & Aga, 1978; Dickey, 2006; Edmonds, 1994).

Because competition in sympatry may select for closely-related species to have more distinct niches in sympatry than in allopatry, we hypothesized that the gut microbiota of *P. vindex* and *P. difformis* would be more dissimilar in sympatry than in allopatry. In addition, we hypothesized that the gut microbiome would be more varied (exhibit greater beta diversity) among populations of the broadly distributed, edaphic-generalist *P. vindex*, compared to populations of the narrowly-distributed, sand- specialist *P. difformis* due to the variation in abiotic conditions across the species’ ranges.

## 2. MATERIALS AND METHODS

### 2.1 Sample collection

We collected *Phanaeus* dung beetles from 17 sampling locations, including 5 allopatric *P. difformis* populations, 6 allopatric *P. vindex* locations, and 6 sympatric *P. vindex* and *P. difformis* populations across Texas, Oklahoma, and Kansas (United States) during May and June 2019 in a manner that minimized temporal autocorrelation (Fig. 1; Table S1). For this study, we considered a site allopatric or sympatric based on the species of beetles we encountered in the field. We identified beetles as *P. vindex* or *P. difformis* using a 30x magnifying hand lens. To differentiate species, we looked for the presence or absence of one or more continuous mid-longitudinal costae on the first or second interstriae of the elytra as well as broad and flat or narrow striae (Dickey, 2006; Edmonds, 1994). We defined a population as all beetles collected within a maximum of 0.5 km^2^, and populations were at least 6 km apart. All sampling was done with proper permission and permitting.

**Figure 1:**
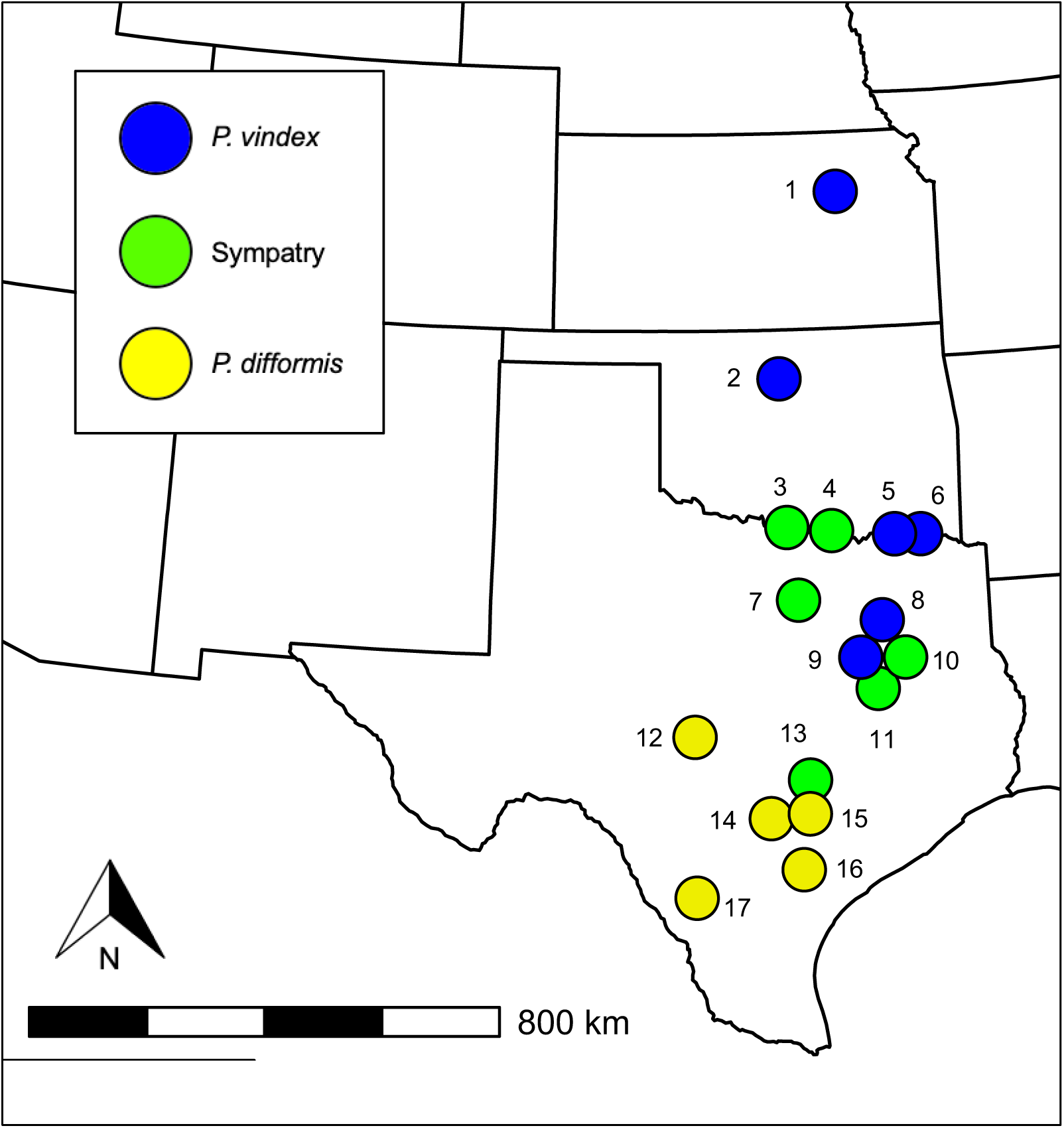
Map showing collection localities of *Phanaeus vindex* and *P. difformis* samples in the states of Kansas, Oklahoma, and Texas (USA). For legibility, points close to each other have been moved slightly. See Table S1 for details about sampling sites.

We used pitfall traps baited with omnivore dung (human or pig) to collect up to twelve females of each locally present *Phanaeus* species at each sampling location. We stored beetles individually in plastic containers and fed them autoclaved cow dung for four days to control for differences in diet consumed prior to capture. We sourced all the dung we fed to the beetles from a single organic cattle pasture that was collected on the same morning, homogenized, sterilized by autoclaving twice, and then stored frozen in small batches until use. We cleaned the plastic containers that we stored live beetles in with a 3% bleach solution between uses. After four days, we euthanized beetles in a vial (one beetle per vial) by submersion in 96% molecular grade ethanol (Fisher BioReagents). We then placed the vials at 4 °C or briefly on ice during transport until further beetle processing.

To examine background soil bacterial and archaeal diversity, we took two soil samples per population, each from one meter away in a random direction from a successful beetle trap. We used flame-sterilized tools to take approximately 0.75 g of soil from a depth of 10 cm. We kept soil samples in 96% molecular grade ethanol at 4 °C or briefly on ice to match beetle storage conditions. High percentages of ethanol (i.e. 95%-100%) are effective at preserving microbial communities in insect guts and soils, among other sample types (Estes et al., 2013; Hammer, Dickerson, & Fierer, 2015; Harry, Gambier, & Garnier-Sillam, 2000).

### 2.2 DNA extractions, 16S library preparation, and MiSeq sequencing

We used the DNeasy PowerSoil Kit (Qiagen, Venlo, Netherlands) to extract DNA from dissected guts and soil samples in a random order to minimize batch effects. Immediately prior to each dissection, we removed the euthanized beetle from its vial of ethanol and weighed it. We dissected out the whole gut of each beetle using flame-sterilized tools on individual pieces of autoclaved aluminum foil. For soil samples, we dried the samples for approximately 35 minutes in a sterilized laminar flow hood on individual sheets of autoclaved aluminum foil. We followed Qiagen’s standard Quick-Start protocol (June 2016), except that we eluted with 50 μL of solution C6 at step 19 and then waited 5 minutes before centrifuging. To characterize the background DNA present in the reagents, we sequenced an extraction kit blank. Although gel electrophoresis post-PCR indicated a very weak band for the extraction blank, NanoDrop spectrophotometry showed that the DNA concentration was too low to be reliably quantified via NanoDrop (<10 ng/ μL). We stored extracted DNA at -20 °C until amplicon library preparation.

Preparation of the 16S rRNA gene amplicon library was performed by the University of Tennessee (UT) Genomics Core in Knoxville, TN, USA. Our protocol followed the 16S Metagenomic Sequencing Library Preparation Workflow published by Illumina Corporation (San Diego, CA, USA), except for three key modifications to the amplicon PCR. First, to amplify the V4 region of the 16S rRNA gene, we used the primers 515 forward and 806 reverse (Caporaso et al., 2011), modified slightly with the addition of two degenerate base pairs (Apprill, McNally, Parsons, & Weber, 2015). The primer sequences of 515F and 806R were GTGYCAGCMGCCGCGGTAA and GGACTACNVGGGTWTCTAAT, respectively. Second, PCR was performed in triplicate, and we reduced each reagent in the amplicon PCR by half and added 0.5 μL of 10 mg/μL bovine serum albumin (Sigma-Aldrich, St. Louis, MO, USA) for a total reaction volume of 13 μL. Finally, the amplicon PCR was carried out in 35 cycles. We ran the amplicon PCR products from each of the triplicate reactions on a 2% agarose gel to verify the amplicon length, then pooled to a volume of 25 μL before performing the index PCR as specified in the Illumina protocol. After completing the index PCR, we pooled the samples to approximately equimolar concentrations using the results from a NanoDrop and confirmed amplicon length and quantity on a Bioanalyzer (Agilent, Santa Clara, CA, USA). We conducted PCR in a UV-sterilized laminar flow hood and included 3 PCR negative controls to verify that minimal DNA contaminants were present in the samples. We were unable to detect PCR negative controls on the agarose gels post PCR, and the DNA concentrations were less than 10 ug/μL when measured via NanoDrop spectrophotometry. To minimize the potential for contamination, we used aerosol barrier pipette tips for all lab work.

Illumina MiSeq sequencing was conducted at the UT Genomics Core facility using a MiSeq Reagent Kit v3 cycle flow cell (500 base pairs) to obtain two 250 bp paired-end reads. We used a loading concentration of 4 pM, and to increase base pair diversity on the flow cell, we spiked in 20% PhiX control DNA (Illumina). We sequenced 125 *P. vindex* samples, 125 *P. difformis* samples, 34 soil samples, 1 extraction blank, and 3 negative PCR controls for a total of 288 samples on the flow cell.

### 2.3 Bioinformatics

We conducted all bioinformatic steps using plugins in QIIME2 (Bolyen et al., 2019), all of which were version 2020.2.0 unless otherwise noted. First, we used cutadapt to remove the 515F and 806R primers from our raw reads (Martin, 2011), allowing an error rate of 0.2. To trim, denoise, dereplicate, and merge our reads, we used DADA2 (Callahan et al., 2016), which resulted in amplicon sequence variants (ASVs) (Yilmaz et al., 2014). We employed DADA2’s default settings and trimmed forward reads at 185 bp and reverse reads at 170 bp. We used scikit-learn version 0.22.1 to train a naïve Bayes taxonomic classifier on the SILVA 128 99% OTUs reference database for our primer region of the 16S gene, and then we used scikit on the trained classifier to assign taxonomy to our sequences. We employed the SATé-enabled phylogenetic placement (SEPP) method (Mirarab, Nguyen, & Warnow, 2012) to place ASVs onto the existing SILVA 128 99% OTUs phylogeny, using the QIIME2 plugin fragment-insertion (Janssen et al., 2018; Eddy, 2011; Matsen, Hoffman, Gallagher, & Stamatakis, 2012; Matsen, Kodner, & Armbrust, 2010). SEPP uses hidden Markov models (Eddy, 2011; Finn, Clements, & Eddy, 2011) trained on the reference tree to cluster sequence fragments more accurately and precisely than do traditional de novo tree-building methods with short sequences such as the 16S gene (Janssen et al., 2018), and is now the method that QIIME2 recommends. We excluded reads that were not present at least two times, ASVs identified as unassigned at the phylum taxonomic level, and chloroplast and mitochondria sequences.

The remainder of the bioinformatics and the statistical analyses were conducted in R (version 4.0.0, R Core Team, 2020). We exported the ASV tables, taxonomy, and the phylogenetic tree from QIIME2, combining them with a metadata table (see next section for details) in R to create a phyloseq object (phyloseq package, version 1.32.0, McMurdie & Holmes, 2013). We obtained the mean ASV abundance for each sample by rarefying 1,000 times to 3,507 and 26,336 reads for gut and soil samples respectively, using rrarefy in the vegan package in R (Oksanen et al., 2019). Using the rarefied gut communities, we constructed a quantitative Jaccard (Ružička) dissimilarity matrix using vegan’s function vegdist and a matrix of weighted UniFrac distances using phyloseq. UniFrac calculates the distance between two samples as the proportion of unshared tree branch lengths out of all tree branch lengths (Lozupone & Knight, 2005). All analyses were repeated for weighted UniFrac and Jaccard dissimilarities.

### 2.4 Metadata sourcing

To understand if sympatry and allopatry explain variation in the gut microbiomes of *P. vindex* and *P. difformis*, we had to account for environmental and geographic/spatial factors that may also cause populations of *Phanaeus* to differ in their gut microbiomes. We utilized the GIS map portal of the National Oceanic and Atmospheric Administration (NOAA) to compute mean monthly temperature, monthly temperature variability (as temperature range), and mean monthly precipitation using data from the years 2009-2018, or the most recent available year prior to 2009 if data from a year between 2009-2018 was unavailable. NOAA measurement stations were no more than 34 km from each sampling location and were on average 20.7 km away. We included an ordinal variable called “Cattle Presence Rank” as an estimate of how many cattle were present at each sampling location. Cattle Presence Rank had three levels: no cows in the beetle collection area or in an area immediately adjacent (“low”), cattle in adjacent area but not present in sampling area (“medium”), or cattle present in sampling area (“high”). To account for the microbial diversity in the beetle’s soil habitat that could influence the beetle gut microbiome, we computed the mean exponential Shannon entropy index for the soil samples taken from each sampling area. Averaging the Shannon indices by sampling site was necessary because one soil sample out of the 34 we sequenced failed (Table S1).

Ecological processes that occur at a variety of spatial scales are important for shaping species interactions (Borcard, Legendre, Avois-Jacquet, & Tuomisto, 2004). Thus, to tease out the effects of spatial autocorrelation on variation in the gut microbiome, we used distance-based Moran’s eigenvector map (dbMEM) models (Borcard & Legendre, 2002; Borcard, Gillet, & Legendre, 2018; Borcard et al., 2004; Dray, Legendre, & Peres-Neto, 2006). The dbMEM approach decomposes spatial relationships among sites, resulting in eigenvectors that describe spatial patterns across the entire range of scales detectable given the coordinates of the samples. We used the function dbmem in the package adespatial (Dray et al., 2020) to create positive and negative dbMEM eigenvalues in a series of three steps. First, we constructed a pairwise Euclidean distance matrix among all sites using geodetic, Cartesian coordinates (in our case, taken from the center of each of our sampling locations). Next, we truncated distances in the matrix based on a threshold value equal to the maximum distance between a pair of neighboring sites. We replaced all pairwise distances larger than the threshold, as well as the matrix diagonals, with the threshold value multiplied by four. Thirdly, we computed a PCoA of the truncated distance matrix, the resulting eigenfunctions describing spatial structure at a range of scales. Prior to testing the explanatory power of the constructed dbMEMs, we removed the effects of linear distance among sites by regressing Jaccard and weighted UniFrac dissimilarity matrices on the X-Y coordinates and then saving the model residuals. We implemented distance-based redundancy analyses (db-RDAs, Legendre & Anderson, 1999) and permutational ANOVAs on dbMEMs with positive and negative eigenvalues separately using the detrended dissimilarity matrices as the response variables. If dbMEM variables were significant, we determined which dbMEMs to include in downstream analyses with the help of forward model selection using vegan function ordiR2step. Finally, we applied the Šidák (1967) correction to P*-*values generated by ordiR2step to account for running multiple tests on positive and negative dbMEMs (Blanchet, Legendre, & Borcard, 2008; Borcard et al., 2018).

We also performed forward model selection separately on environmental variables (i.e. mean temperature, temperature variation, mean precipitation, cattle presence, and soil habitat microbial biodiversity) using vegan function ordiR2step (Table S2). Following these model selections, we created two separate R objects from the combined geographic variables (significant MEMs and X,Y coordinates because we detected a linear trend in our data) and the significant environmental variables. Performing multiple rounds of model selection before hypothesis testing was necessary to avoid saturating our models with our many initial candidate predictors. In addition, it allowed us to understand how environmental, geographic, and biotic variables may be jointly influencing patterns of gut microbial variation via variation partitioning.

### 2.5 Examining the drivers of variation in the gut microbiome

We performed variation partitioning using the vegan function varpart to understand the explanatory power of our geographic variables, environmental variables, range overlap, and *Phanaeus* species in relation to variation in the gut microbiome. Variation partitioning runs independent redundancy analyses on the response variables and each matrix of explanatory variables, and then computes the adjusted *R^2^* of each fraction via subtraction (Borcard et al., 1992; Peres-Neto, Legendre, Dray, & Borcard, 2006). Therefore, this technique allowed us to understand the unique and overlapping contributions of all of the types of variables in our models, including those that were correlated.

To test our hypothesis that the gut microbiomes of *P. vindex* and *P. difformis* would be more similar in allopatry than in sympatry, we implemented db-RDAs using the Jaccard and the weighted UniFrac dissimilarity matrices as the response variables. Prior to implementing db-RDAs, we performed global model selection using ordiR2step on the environmental and geographic variables used in variation partitioning, “patry” (sympatry or allopatry), species, and beetle mass (Table S3). Based on these model selections, we included X,Y coordinates (i.e. longitude and latitude), five negative dbMEMs (which account for spatial relationships), the mean Shannon entropy of soil samples, cattle presence, species, range overlap, and the interaction between species and range overlap as predictors in our quantitative Jaccard db-RDA. Despite forward selection identifying mean precipitation as a significant variable, we did not use it in our Jaccard model because it had a variance inflation factor (VIF) of over 20, presumably because of a high correlation with longitude (Pearson correlation coefficient = .901). For our model using the weighted UniFrac dissimilarity as the response variable, we included the Y coordinates (i.e. latitude), a single significant negative dbMEM, mean precipitation, cattle abundance, species, range overlap, and the interaction between species and range overlap as predictors. Other than the interaction term and its variables, all variables had a VIF of 5 or lower. Testing the significance of the interaction between species and range overlap allowed us to explicitly consider if the gut microbiomes of *P. difformis* and *P. vindex* differentially depend on the presence of the other species. We investigated the significance of our models’ constraints using the vegan function anova.cca with 99,999 permutations. Finally, to separate out the proportions of the variation explained by each predictor, we again performed variation partitioning using the same predictors that were in the db-RDAs.

### 2.6 Testing differences in beta diversity among populations of *P. vindex* and *P. difformis*

We hypothesized that the gut microbiome of the broadly-distributed, cosmopolitan *P. vindex* would exhibit greater beta diversity than that of the relatively narrowly-distributed, specialist *P. difformis*. Beta diversity describes the variation in species composition among sampling units for a geographic area of interest (Whittaker, 1960). One way of quantifying beta diversity is as the average distance between sampling units and their group centroid in multivariate space, i.e. multivariate dispersion (Anderson, Ellingsen, & McArdle, 2006). However, to obtain beta diversities representing variation in the gut microbiome among populations of *P. vindex* and *P. difformis*, we had to account for uneven sample size, spatial relationships among our sample sites, and other random factors including cattle abundance and soil biodiversity that could influence the gut microbiome from one site to the next. To do this, we first created mean ASV tables for each population of *P. vindex* and *P. difformis,* and then obtained pairwise weighted UniFrac and quantitative Jaccard dissimilarities among them. Next, in the same way as described above, we identified dbMEMs and random environmental variables that were important for our models using forward selection. Finally, we regressed Jaccard and weighted UniFrac dissimilarities on significant spatial and environmental predictors. To calculate the multivariate dispersions of *P. vindex* and *P. difformis,* we input the residuals of these models into the function betadisper in vegan. We tested the null hypothesis of no difference between the beta diversities of *P. vindex* and *P. difformis* gut communities using function permutest (99,999 permutations).

### 2.7 Indicator species analyses

We used two indicator species analyses to identify potentially important ASVs in the beetles’ guts (De Cáceres & Legendre, 2009; De Caceres, Legendre, & Moretti, 2010), including one analysis to identify gut microbes that were characteristic of *P. vindex* and *P. difformis* in sympatry or in allopatry, and a second analysis to examine gut microbes that were characteristic of the abundance of cattle in the sampling locations. The first analysis identified ASVs with different affinities for *P. vindex* and *P. difformis* in sympatry and in allopatry, both species in sympatry, and both species in allopatry. The second analysis identified ASVs associated with “low”, “medium”, and “high” abundances of cattle in the beetles’ environment. To perform the indicator species analyses, we estimated the group equalized, point-biserial correlation coefficients, *r_g_*, between ASVs and groups described above and the significance of these associations using 99,999 permutations with the function multipatt in version 1.7.9 of the package indicspecies (De Cáceres & Legendre, 2009). The point-biserial correlation coefficient takes into account the abundance of each ASV in the group, as well as whether or not it occurs in the other groups under consideration. We then calculated the relative abundances of each indicator species in each sample grouping.

## 3. RESULTS

### 3.1 Characterization of *Phanaeus* gut microbial communities

Our MiSeq sequencing run yielded 11,130,657 raw sequences. For all samples, the median number of sequences per sample was 37,458, and for beetle gut and soil samples, the median number of sequences per sample was 33,721 and 62,157, respectively. After trimming, merging, and chimera removal with DADA2, 6,442,702 sequences remained representing 33,488 unique ASVs. Of these, we retained 23,641 bacterial or archaeal ASVs that were assigned to at least the phylum level and that appeared at least twice in our dataset. This included a mean of 18,310 sequences per gut sample and a mean of 40,699 sequences per soil sample. Rarefaction curves indicated that a sampling depth of 3,500 and 26,336 for gut and soil samples, respectively, was more than adequate to capture the full microbial richness of the communities (Fig. S4). After rarefying, we retained 110 *P. difformis*, 89 *P. vindex*, and 33 soil samples. Because we took the mean communities of 1,000 rarefactions, we retained all 23,641 original ASVs, which included 855 ASVs in *P. vindex* guts, 907 ASVs in *P.* difformis guts, and 22,365 ASVs in soil samples.

*P. vindex* and *P. difformis* gut communities were dominated by the same bacterial phyla and families; however, the relative abundances of the top phyla (Fig. 2) and families (Fig. 3) differed between the beetle species. The most abundant phyla included Firmicutes (*P. vindex:* 39%, *P. difformis:* 43%), Proteobacteria (*P. vindex*: 37%, *P. difformis:* 31%), Bacteroidetes (*P. vindex*: 16%, *P. difformis*: 19%), and Actinobacteria (*P. vindex*: 5.%, *P. difformis:* 7%). In addition, 1% of the ASVs found in *P. vindex* belonged to the phylum Fusobacteria. No other phylum characterized more than 2% of the ASVs in either *Phanaeus* species. The five most abundant bacterial families were Enterococcaceae (*P. vindex*: 24%, *P. difformis*: 28%), Moraxellaceae (*P. vindex*: 18%, *P. difformis*: 10%), Porphyromonadaceae (*P. vindex*: 11%, *P. difformis*: 16%), Enterobacteriaceae (*P. vindex*: 14%, *P. difformis*: 13%, and Planococcaceae (*P. vindex*: 9%, *P. difformis*: 9%). All remaining families characterized fewer than 5% of the ASVs we found in *P. vindex* and in *P. difformis*. Although our primers were designed to detect archaeal and bacterial 16S rDNA, Archaea represented only 0.003% of reads in both *P. difformis* and *P. vindex*.

**Figure 2:**
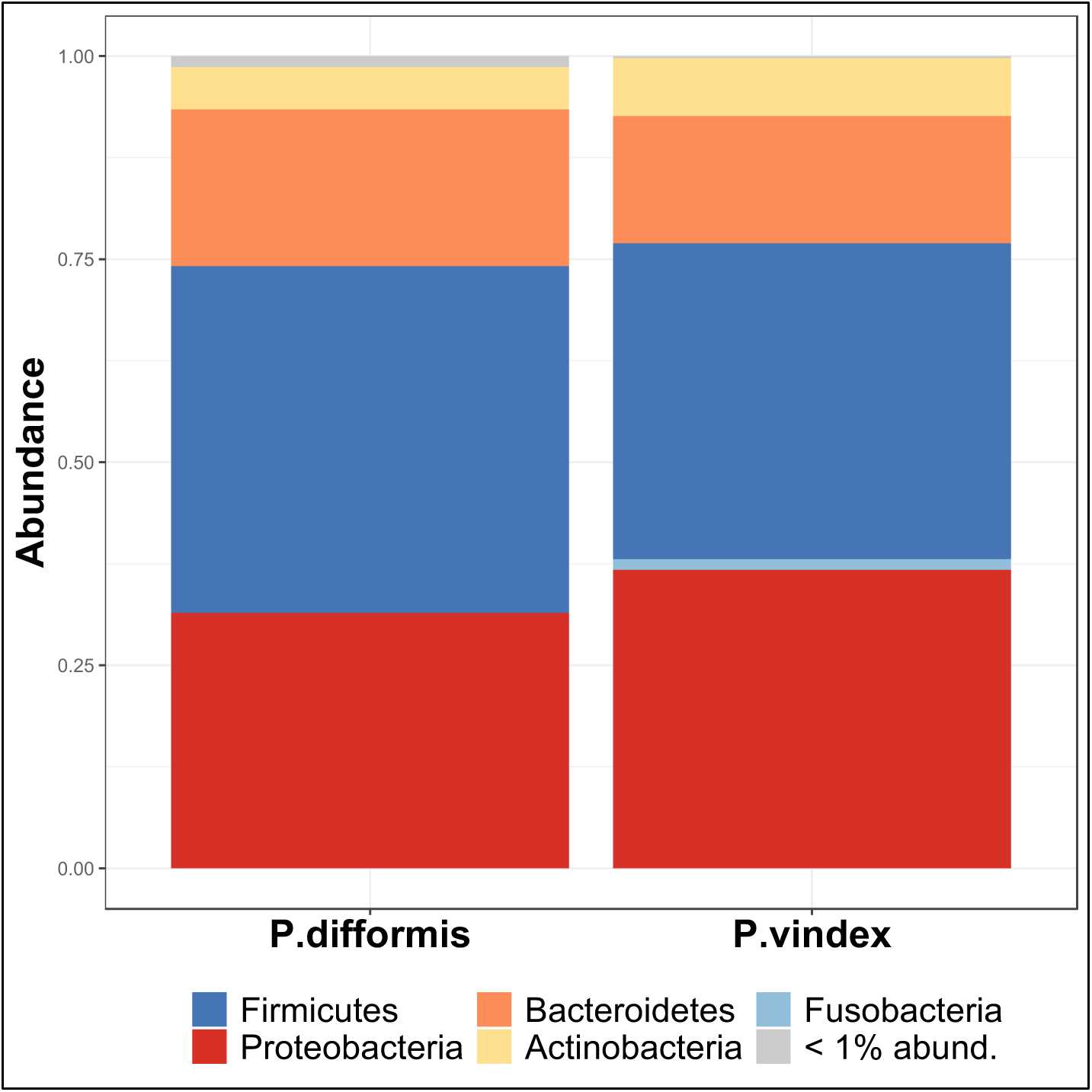
Relative abundance of taxonomic phyla present in the guts of *Phanaeus difformis* and *P. vindex* samples. Taxonomic groups representing less than 1% abundance have been combined.

**Figure 3:**
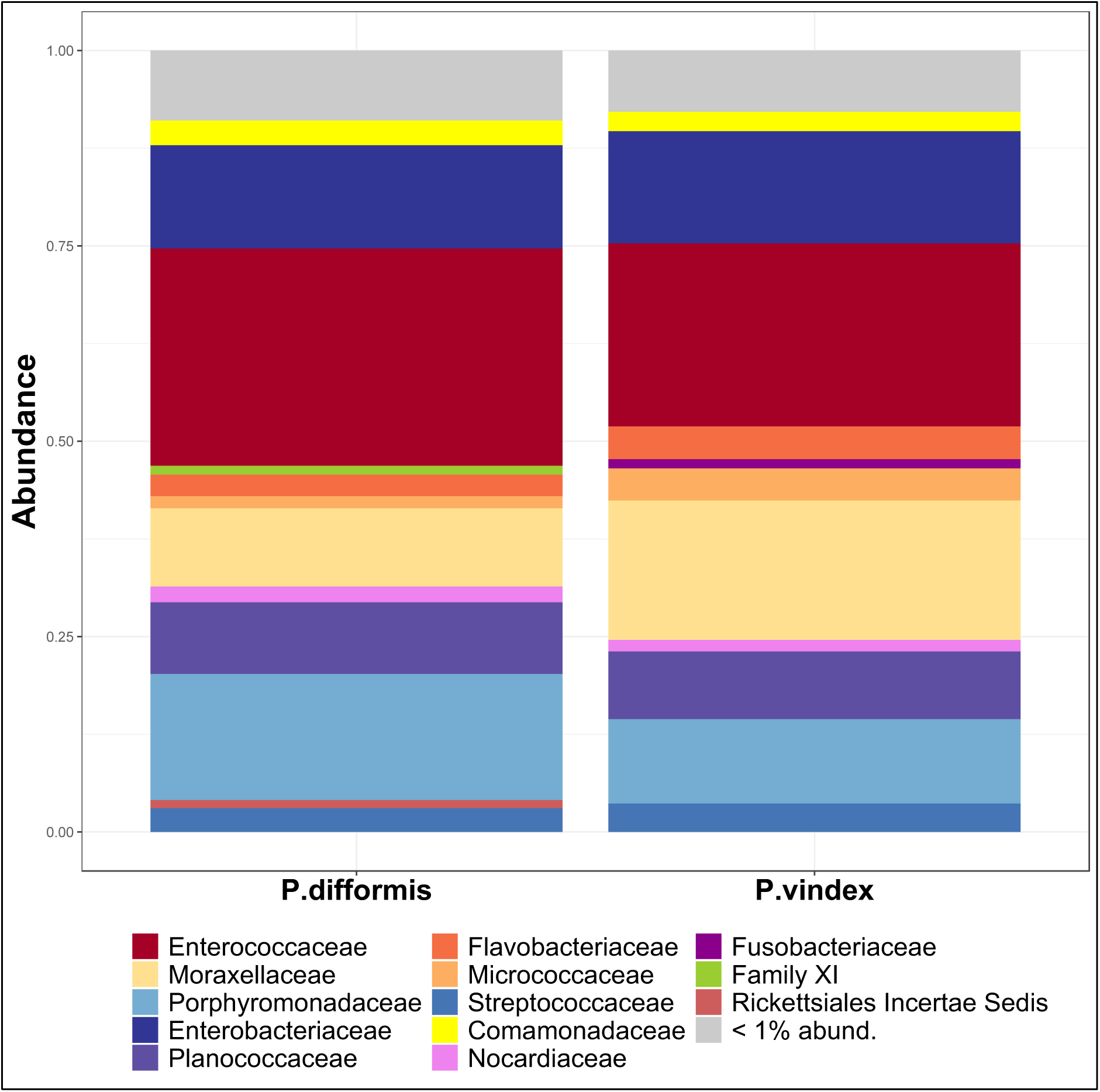
Relative abundance of taxonomic families present in the guts of *Phanaeus difformis* and *P. vindex* samples. Taxonomic groups representing less than 1% abundance have been combined.

We performed two indicator species analyses to identify potentially important ASVs in the beetles’ guts, including one based on sympatry or allopatry of the beetles (Table S5) and one based on the abundance of cattle in the beetles’ environment (Table S6). We found 21 indicator species for allopatric *P. difformis*, 11 for sympatric *P. difformis*, 20 for allopatric *P. vindex*, and 15 for sympatric *P. vindex*. For low, medium, and high abundance of cattle, we identified 12, 40, and 42 ASVs, respectively. In both analyses, the identified indicator species represented between 10 and 20% of the total abundance of ASVs; however, the overwhelming majority of indicator species comprised less than 1% of a group’s total relative abundance. Notable indicator taxa in the first analysis that were both highly correlated and highly abundant included an unidentified ASV in Planococcaceae for allopatric *P. difformis* (*r_g_* = 0.267, abundance = 2%), two *Vagococcus spp.* in sympatric *P. difformis* samples (*r_g_* = 0.363 and 0.202; abundances = 13% and 4%), *Acinetobacter sp*. (*r_g_* = 0.278, abundance = 7%) and an unknown ASV within Enterobacteriaceae (*r_g_* = 0.318, abundance = 5%) for allopatric *P. vindex*, and an ASV within the genus *Glutamicibacter* (*r_g_* = 0.311, abundance = 1%) for sympatric *P. vindex* samples. Importantly, our second indicator analysis revealed that the abundance of several indicator taxa showed a positive relationship with cattle abundance, suggesting that the ASVs are acquired via the consumption of cattle dung. The most abundant of these taxa included *Vagococcus sp.* (high: 9%, medium: 5%, low: 3%)*, Gordonia sp*.(high: 3%, medium: 1%, low: 1%), and *Acinetobacter sp.* (high: 2%, medium: 1%, low: 1%). Different ASVs within the genus *Dysgonomonas* were identified as indicator taxa for all four groups of *P. vindex* and *P. difformis* samples as well as high and medium cattle abundance. Interestingly, three distinct ASVs that are known endosymbionts of the dung beetle *Onthophagus taurus* were found in the sympatric *P. difformis* group (a Rhodobacteraceae ASV), the low cattle abundance group (a Pasteurellaceae ASV) and the high cattle abundance group (a different Pasteurellaceae ASV). Finally, we found up to three indicator ASVs for all combinations of *P. vindex* and *P. difformis* samples in sympatry or in allopatry, except for the combination of *P. vindex* and *P. difformis* sympatric samples. No ASVs had a high affinity for multiple levels of cattle abundance.

### 3.2 Contributions of range overlap, *Phanaeus* species, and environmental and spatial variables to gut microbiome structure

Based on quantitative Jaccard dissimilarities, which are phylogenetically-blind but take into account ASV richness and abundance, differences in the gut microbiome among individual beetles were high, with dissimilarity values of between .8 and .95 across all sympatric and allopatric groups (Fig. 4a). In contrast, phylogenetically-informed weighted UniFrac dissimilarity values were much lower but still very uniform across categories, ranging from .2 to .4 (Fig. 4b). The difference between quantitative Jaccard and weighted UniFrac values indicates that although ASVs are highly variable among individuals, these ASVs are closely related, and thus are likely ecologically similar, too. Allopatric *P. vindex* individuals appeared to be more different from one another than individuals within the other categories, particularly in the weighted UniFrac analysis, but this difference was small. There appeared to be no difference in the dissimilarity values between allopatric *P. vindex* and *P.* difformis, and sympatric *P. vindex* and *P. difformis*.

**Figure 4:**
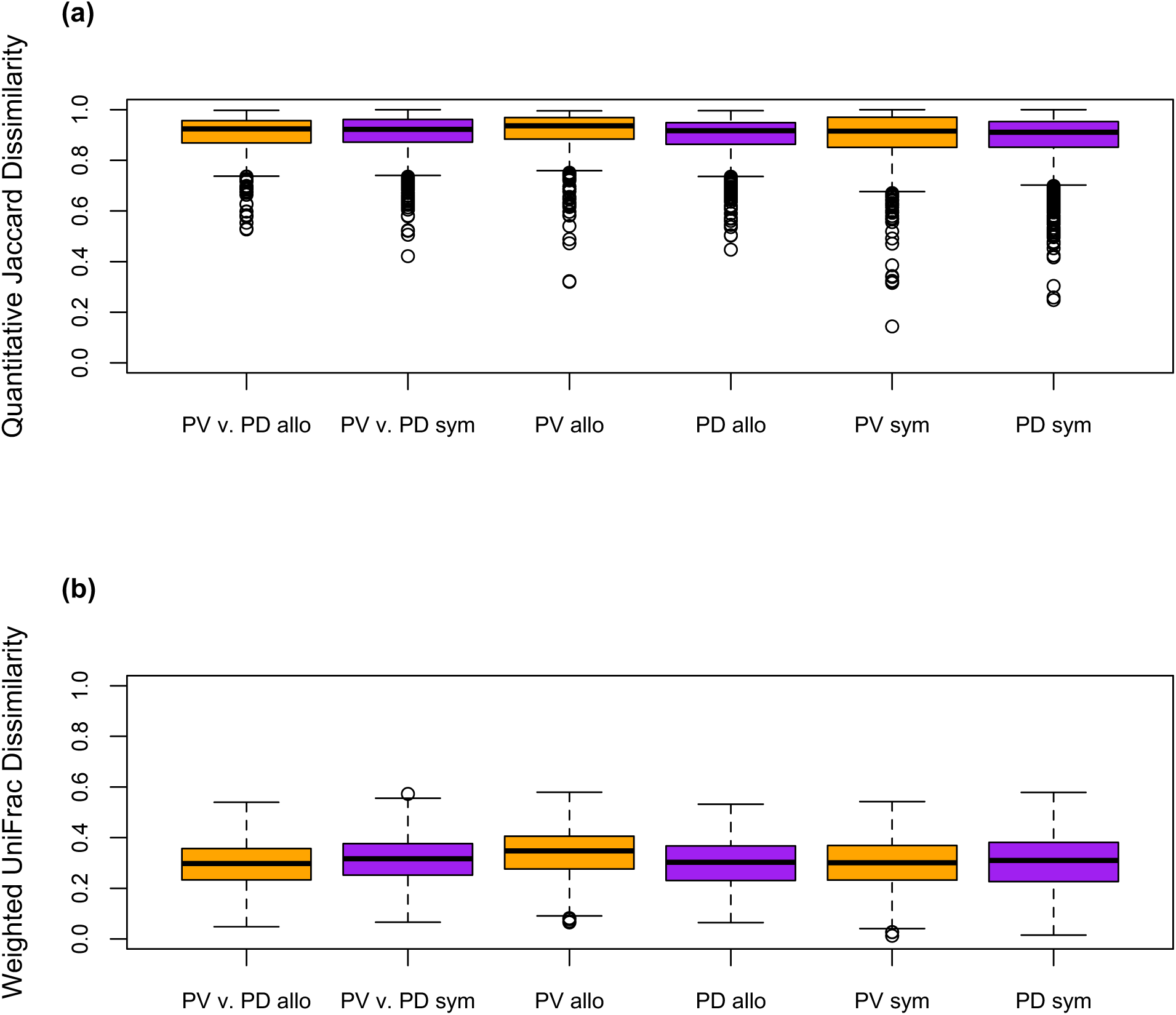
Boxplots comparing (a) quantitative Jaccard dissimilarities and (b) weighed UniFrac distances among different populations of *Phanaeus vindex* and *P. difformis.* Orange boxes show allopatric comparisons, while purple boxes indicate sympatric comparisons. From left to right, comparisons are among allopatric individuals of *P. vindex* and *P. difformis*, among sympatric individuals of *P. vindex* and *P. difformis*, among allopatric *P. vindex* individuals, among allopatric *P. difformis* individuals, among sympatric *P. vindex* individuals, and among sympatric *P. difformis* individuals. Horizontal lines in each boxplot indicate the median dissimilarity value.

Because the simple comparisons of dissimilarity values do not take into account other likely important sources of variation in the beetles’ environments, we used variation partitioning to explore how the environmental, geographic (significant dbMEMs and X-Y coordinates), and biotic variables (“patry” and *Phanaeus* species) contributed to the overall variation of the gut microbiome data. In our Jaccard model, variation in the gut microbiome was explained by geographic variables (2.8%), followed by environmental variables (1.1%), species (1.0%), and sympatry or allopatry (.3%) (Fig. S7a). In our weighted UniFrac model, variation in the gut microbiome was accounted for by geographic variables (1.1%), environmental variables (1.8%), species identity (1.0%), and range overlap (1.1%) (Fig. S7b). Positive dbMEM eigenvalues did not explain a significant amount of the variation in our detrended dissimilarity matrices (Jaccard: *p* = .70, weighted UniFrac: *p* = .13); thus, they were not included in analyses. More information on model selection is given in Table S2.

Permutational ANOVAs revealed that different variables explained weighted UniFrac and Jaccard dissimilarities among gut samples. Surprisingly, geographic predictors (*p* < .01), mean precipitation (*p* < .001), and cattle abundance (*p* < .05) were the only significant explanatory variables in our weighted UniFrac model (Fig. 5). However, the interaction term between *Phanaeus* species and range overlap was significant (*p* < .01) in our model with the Jaccard dissimilarity matrix, as were geographic variables (*p* < .0001) and cattle abundance (*p* < .001) (Fig. 6). Variation partitioning analyses based on the predictors used in the quantitative Jaccard db-RDA model indicated that geography, environmental variables, species, and range overlap respectfully contributed about 2.8%, 0.8%, 1.0%, and 0.4% to the overall variation in the data. Despite our attempt to reduce collinearity, 0.3% of the variation was explained jointly by geography and the environmental variables. For the predictors used in the weighted UniFrac model db-RDA, variance partitioning showed that 2.4%, 1.8%, 1.1%, and 0.6% were explained by geography, environmental variables, species, and range overlap, respectively.

**Figure 5:**
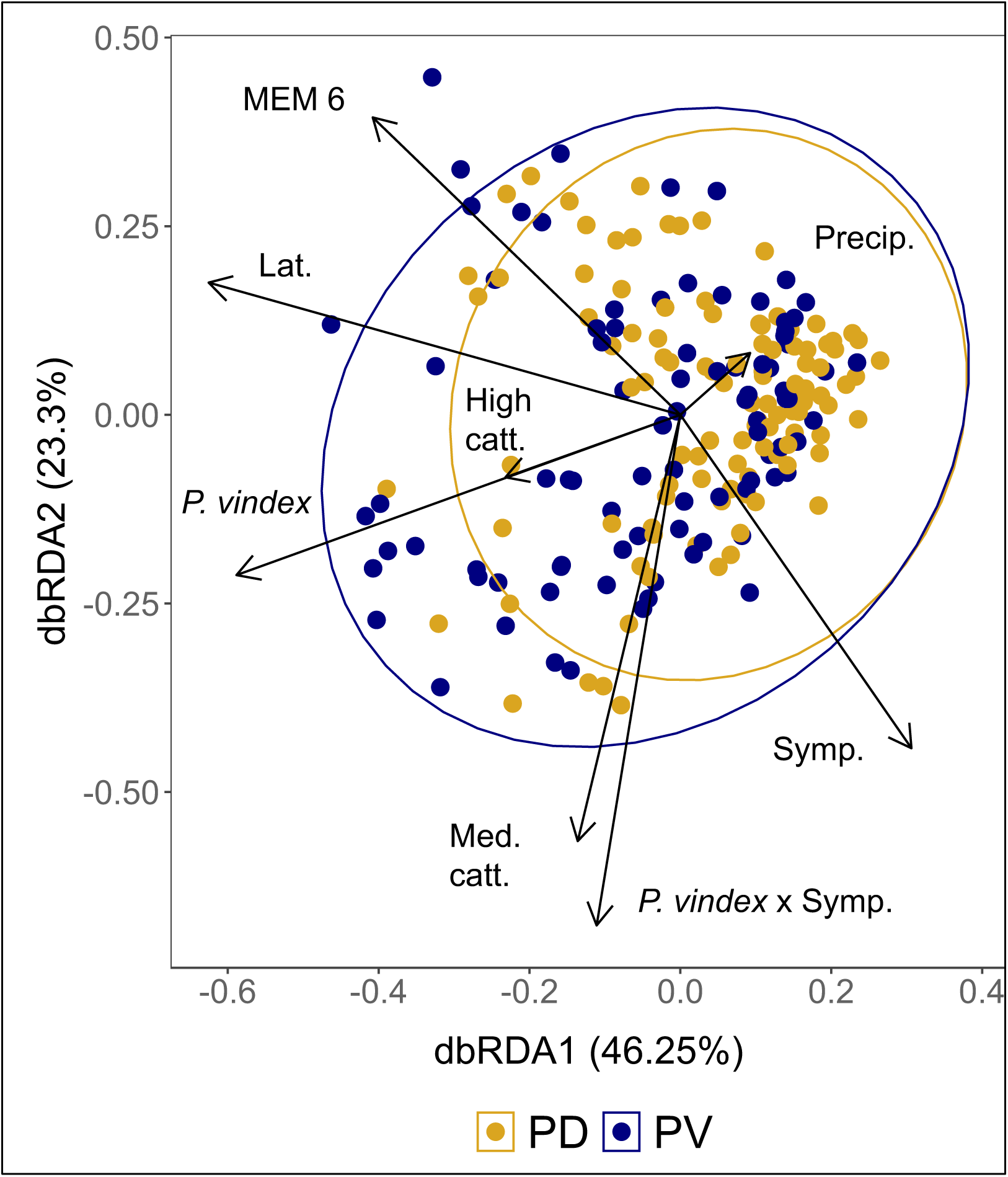
Plot of distance-based redundancy analysis (db-RDA) based on weighted UniFrac dissimilarity matrix of *Phanaeus difformis* and *P. vindex* samples. Points on the plot indicate individual *Phanaeus* beetles. Ellipses represent 95% confidence intervals around centroids of each *Phanaeus* species. All constrained axes together explain 10.00% of the total variation in multivariate space. Arrows indicate the strength of correlation of variables with dbRDA axes 1 and 2. Numerical variables constraining the ordination include one negative distance-based Moran’s eigenvector mapping eigenvalues (MEM 6), latitude (Lat.), and precipitation of the sampling location (Precip.). Constraining factor variables include cattle presence in the sampling area (low, medium, or high), *Phanaeus* species, range overlap (i.e. sympatry or allopatry) and the interaction between *Phanaeus* species and range overlap (*P. vindex* x Symp.). Factor variables are automatically dummy coded; thus, the factor level coded as the intercept does not have a corresponding axis. A permutational ANOVA (99,999 permutations) showed that geographic predictors (i.e. MEM 6 and latitude), precipitation, and the abundance of cattle at the sampling location were significant predictors in the model.

**Figure 6:**
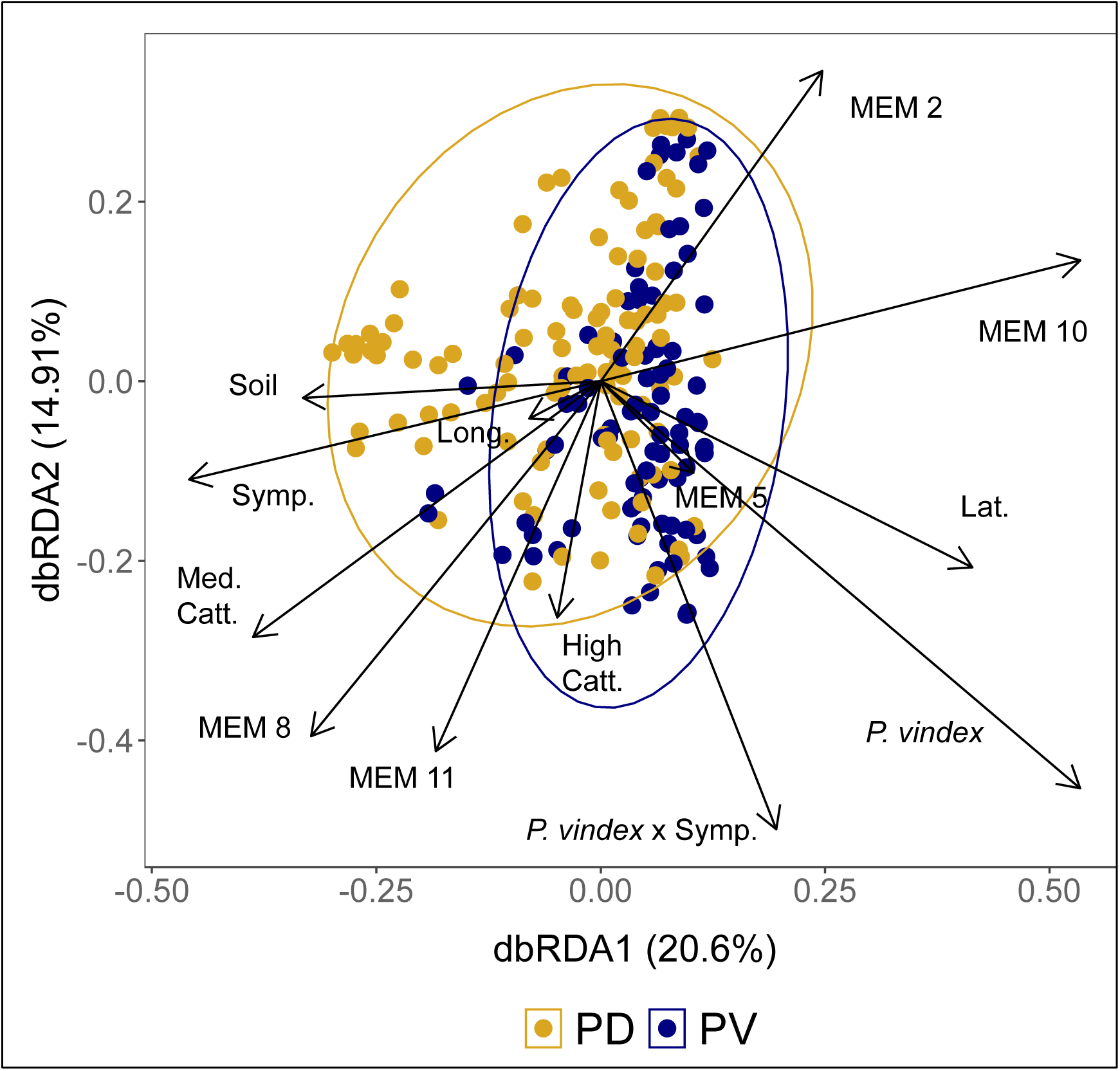
Plot of distance-based redundancy analysis (db-RDA) based on the quantitative Jaccard dissimilarity matrix of *Phanaeus difformis* and *P. vindex* samples. Points on the plot indicate individual *Phanaeus* beetles. Ellipses represent 95% confidence intervals around centroids of each *Phanaeus* species. All constrained axes together explain 12.27% of the total variation in multivariate space. Arrows indicate the strength of correlation of variables with dbRDA axes 1 and 2. Numerical variables constraining the ordination include five negative distance-based Moran’s eigenvector mapping eigenvalues (MEMs 2, 5, 8, 10, and 11), latitude (Lat.), longitude (Long), and the mean Shannon index of soil samples taken from each sampling location (Soil). Constraining factor variables include cattle presence in the sampling area (low, medium, or high), *Phanaeus* species, range overlap (i.e. sympatry or allopatry) and the interaction between *Phanaeus* species and range overlap (*P. vindex* x Symp). Factor variables are automatically dummy coded; thus, the factor level coded as the intercept does not have a corresponding axis. A permutational ANOVA (99,999 permutations) indicated that geographic variables (i.e. dbMEMs, latitude, and longitude), cattle abundance, and the interaction effect between *P. vindex* and *P. difformis* were significant in the model.

To understand the significant interaction between *Phanaeus* species and range overlap in our Jaccard model, we ran this model another three times on Jaccard dissimilarity matrices separated by *P. vindex* and *P. difformis* and on a Jaccard dissimilarity matrix based on only sympatric samples. For these models, we did not include the dbMEMs as predictors because they were constructed using all coordinates, including those from sites where only one *Phanaeus* species was collected. Interestingly, we found that range overlap explained a significant amount of the variation in both the *P. vindex* and the *P. difformis* models (*P. vindex* model: *p* < .01; *P. difformis* model: *p* < .05), while *Phanaeus* species identity was significant in the sympatry model: *p* < .0001). All models were also explained by geographic variables (i.e. X,Y coordinates) (*P. vindex*: *p* < .001; *P. difformis*: *p* < .001; sympatry model: *p* < .0001) and cattle abundance (*P. vindex* model: *p* < .01; *P. difformis* and sympatry models: *p* < .001). However, the mean Shannon index of the soil samples was only significant in the sympatry model (*p* < .01).

### 3.4 Comparisons of the beta diversity of *P. vindex* and *P. difformis* populations

Prior to comparing beta diversities, we removed the effects of random environmental variables (i.e. mean soil Shannon diversity and cattle abundance) and geographic distance (no positive or negative dbMEMs were significant) that could cause random differences among populations of *P. vindex* and *P. difformis*. We did not detect a difference in the beta diversities of *P. vindex* and *P. difformis* for Jaccard (*p* = .39; Fig. 7) or weighed UniFrac models (*p* = .20; Fig. S8).

**Figure 7:**
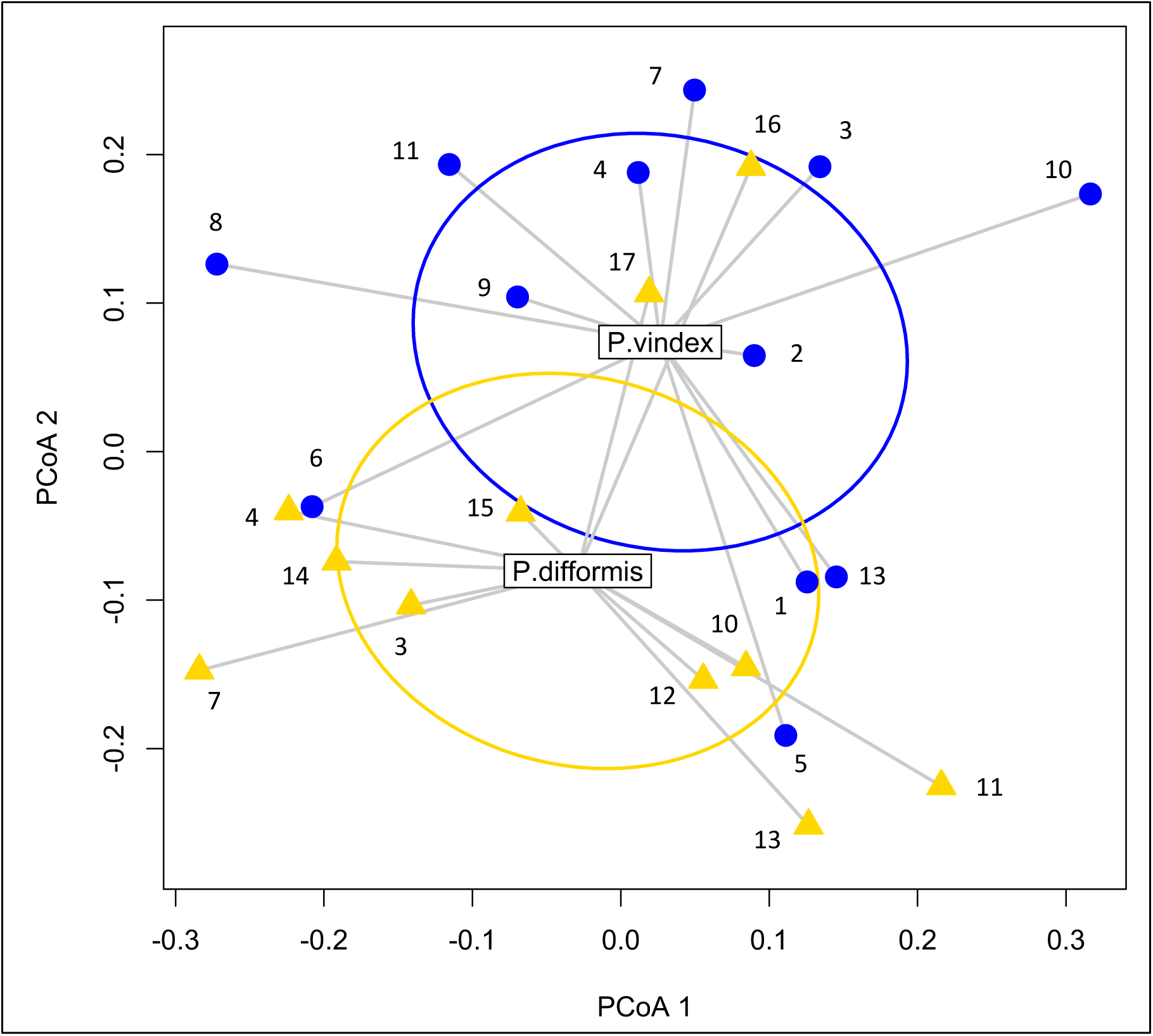
Beta diversity of *Phanaeus vindex* and *P. difformis* represented as multivariate dispersions based on Jaccard dissimilarity dissimilarities. Populations of *P. vindex* are represented by blue dots; gold triangles indicate populations of *P. difformis*. Numbers represent different populations by location (see Table S1 for metadata associated with each population). The words “*P. vindex*” and “*P. difformis*” are positioned in the species’ respective centroids in multivariate space. Ellipses represent one standard deviation away from the centroid of *P. vindex* and *P. difformis*.

## 4. DISCUSSION

### 4.1 Summary of results

In this study, we sampled *Phanaeus vindex* and *P. difformis* dung beetles across their allopatric and sympatric ranges to understand the factors influencing variation in the gut microbiome. We did not find that the gut microbiomes of *P. vindex* and *P. difformis* were more similar in allopatry than in sympatry as we expected. However, our analyses based on Jaccard dissimilarities indicated that the gut microbiomes of *P. vindex* and *P. difformis* respond differently to sympatric and allopatric conditions, consistent with the expectations of character displacement. In addition, we hypothesized that because *P. vindex* is found in more diverse habitats and has a larger geographic range than *P. difformis*, its gut microbiome would vary more among populations. While it appears that *P. vindex* has greater beta diversity than does *P. difformis* in our analysis based on weighted UniFrac dissimilarities (Fig. S8), the results were not statistically significant (*p* = .20). However, because we controlled for the fact that distances among populations were not uniform, we may have been overly conservative in our assessment of beta diversity. Specifically, by removing the effect of spatial structuring in our data, we may have also factored out ecologically-relevant variation in the gut microbiome of widely-distributed *P. vindex*, whose populations were also more spaced out on the landscape that those of *P. difformis*. Alternatively, we may have simply needed more populations of both species. Overall, geographic distance among populations, environmental variables, and local dung sources were the most important for shaping the gut microbiomes of *P. difformis* and *P. vindex*. Finally, taxa identified as highly abundant or as indicator species echo those found by other dung beetle researchers, suggesting candidate taxa for the core vertically-transmitted gut microbiome of Scarabaeinae dung beetles. We discuss our findings and their implications below.

### 4.2 Evidence of character displacement in sympatry

Surprisingly, our db-RDAs and subsequent ANOVAs on the full gut microbiome dataset suggested that *Phanaeus* species identity and range overlap were poor predictors of overall variation in the gut microbiome (Figs. 5 and 6). However, in our models based on quantitative Jaccard dissimilarities, we found a significant interaction effect between *Phanaeus* species and range overlap. This indicates that the gut microbiome of one of the *Phanaeus* species shifts more from allopatry to sympatry than does the gut microbiome of its sister species. When we performed additional analyses on separated *P. vindex* and *P. difformis* Jaccard dissimilarities to disentangle this interaction effect, we found that the gut microbiomes of both *Phanaeus* species differed based on range overlap. While we cannot validate it statistically, it appears that this trend is stronger among *P. vindex* samples (Fig. 8) than among *P. difformis* samples (Fig. 9), despite the fact that many of our allopatric *P. vindex* locations were geographically very close to where we encountered sympatric *Phanaeus* populations (Fig. 1, Table S1).

**Figure 8:**
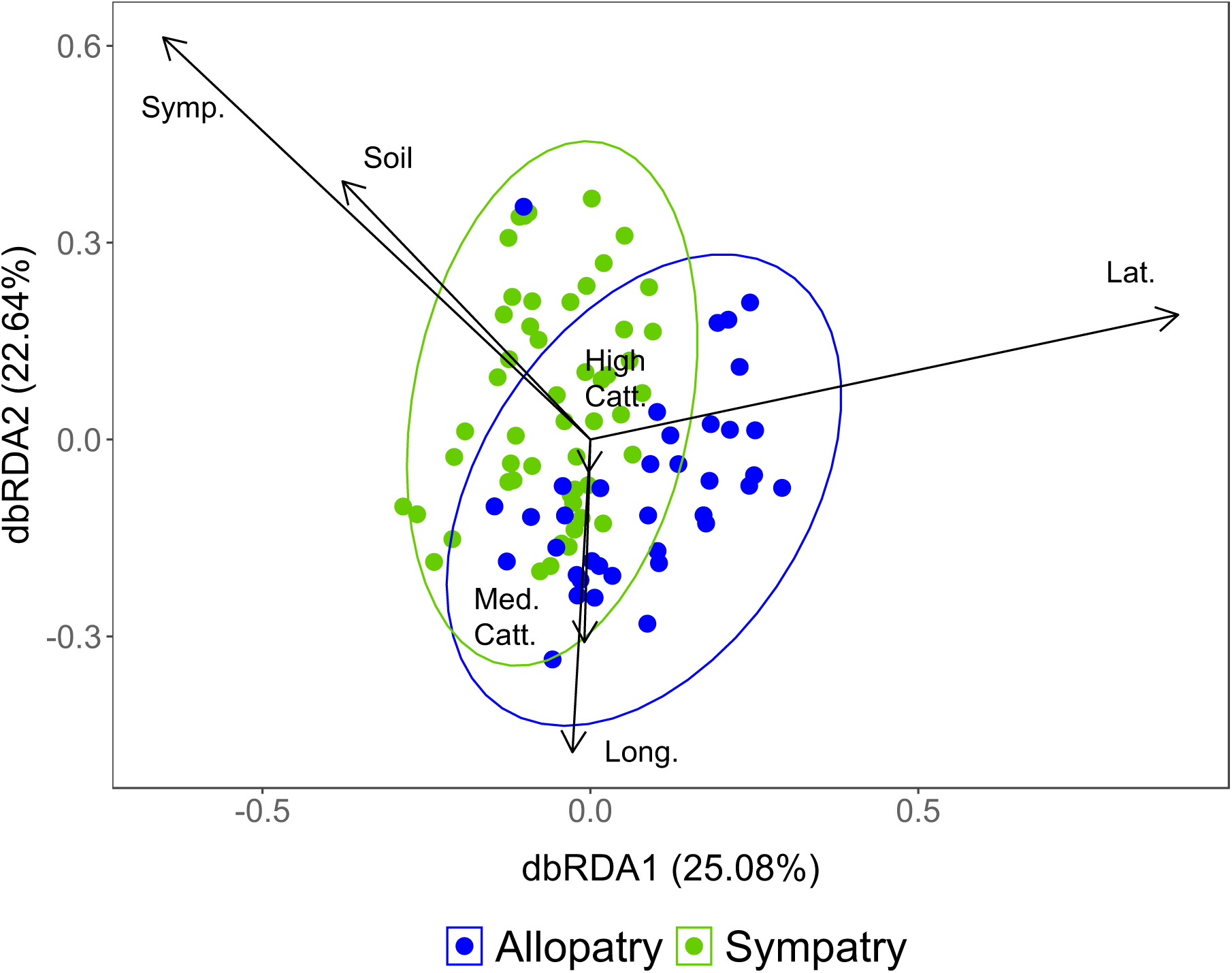
Plot of distance-based redundancy analysis (db-RDA) based on the quantitative Jaccard dissimilarity among *Phanaeus vindex* samples in sympatry and in allopatry. Points on the plot indicate individual *P. vindex* beetles. Ellipses represent 95% confidence intervals around centroids of *P. vindex* in allopatry and in sympatry with *P. difformis*. All constrained axes together explain 10.89% of the total variation in multivariate space. Arrows indicate the strength of correlation of variables with dbRDA axes 1 and 2. Numerical variables constraining the ordination latitude (Lat), longitude (Long.), and the mean Shannon index of soil samples taken from the sampling areas (Soil). Constraining factor variables include cattle presence in the sampling area (low, medium, or high), and range overlap (i.e. sympatry or allopatry). Factor variables were dummy coded; thus, the factor level coded as the intercept does not have a corresponding axis. All variables shown were significant following ANOVAs (99,999 permutations).

**Figure 9:**
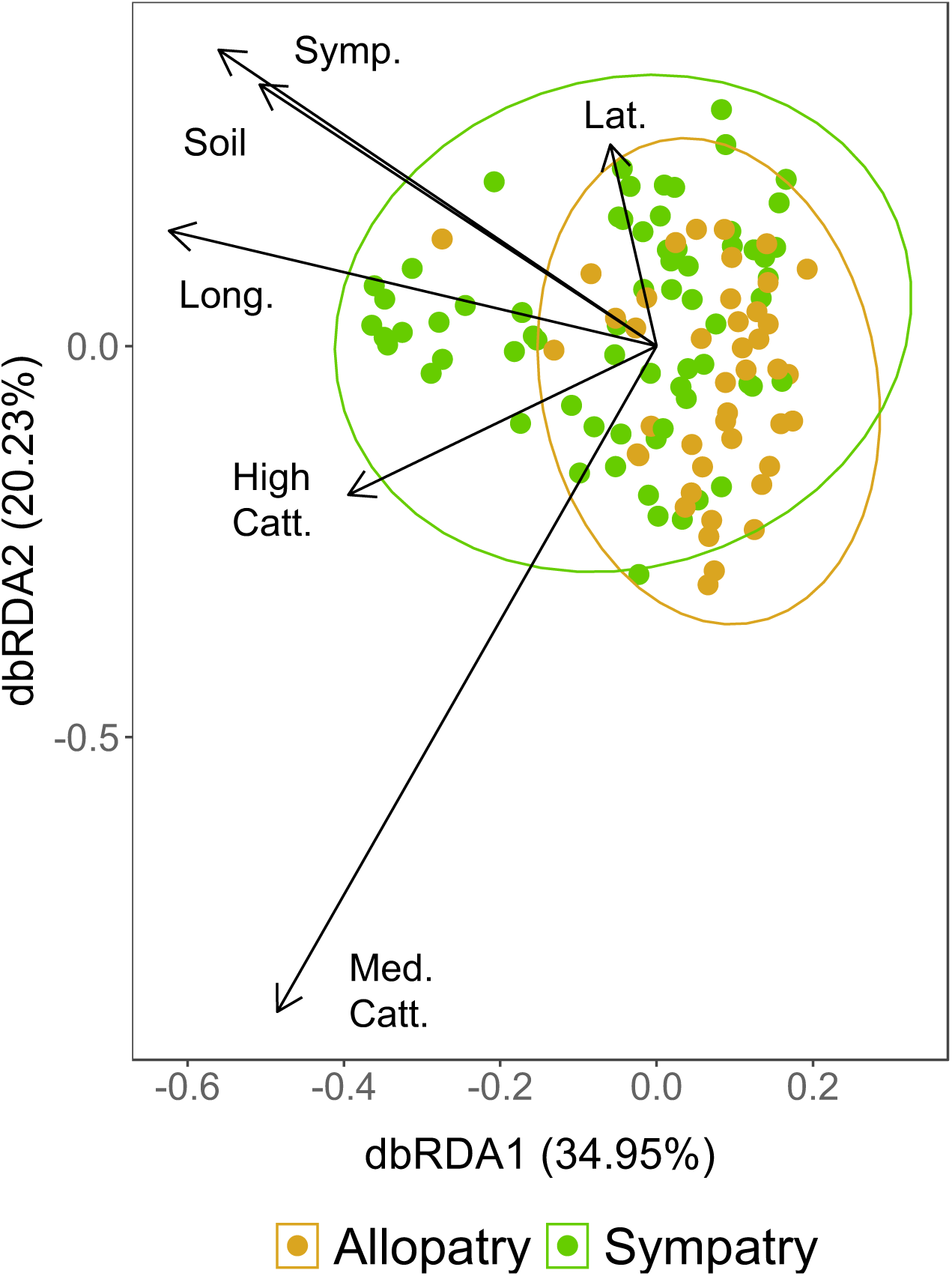
Plot of distance-based redundancy analysis (db-RDA) based on the quantitative Jaccard dissimilarity among *Phanaeus difformis* samples in sympatry and in allopatry. Points on the plot indicate individual *P. difformis* beetles. Ellipses represent 95% confidence intervals around centroids of *P. difformis* in allopatry and in sympatry with *P. vindex*. All constrained axes together explain 9.47% of the total variation in multivariate space. Arrows indicate the strength of correlation of variables with dbRDA axes 1 and 2. Numerical variables constraining the ordination latitude (Lat), longitude (Long.), and the mean Shannon index of soil samples taken from the sampling areas (Soil). Constraining factor variables include cattle presence in the sampling area (low, medium, or high), and range overlap (i.e. sympatry or allopatry). Factor variables were dummy coded; thus, the factor level coded as the intercept does not have a corresponding axis. All variables shown were significant following ANOVAs (99,999 permutations).

Sympatric *Phanaeus* compete fiercely for dung and space to bury brood balls (Price & May, 2009), and thus, ecological theory suggests that their co-existence is predicated on a change in a trait that facilitates niche partitioning (e.g. Brown & Wilson, 1956). We speculate that shifts in the gut microbiome in sympatry may be a form of character displacement which reduces competition between *P. vindex* and *P. difformis* in sympatry and allows for co-existence.

A distinct gut microbiome may aid niche partitioning by facilitating the co-existence of *P. vindex* and *P. difformis* on sandy soils. As *P. difformis* is a sand specialist, we found all of the sympatric populations on sandy soils. *P. difformis* has an additional point on its front tibiae compared to *P. vindex* (Dickey, 2006; Edmonds, 1994), potentially giving it a competitive advantage over *P. vindex* in displacing sandy soil and burying brood balls where the two species co-occur. To reduce competition for space below the dung pat in sympatry, *P. vindex* may bury its brood balls more shallowly in the soil profile than *P. difformis*, a strategy found among sympatric *Onthophagus* dung beetles (Macagno, Moczek, & Pizzo, 2016). Brood balls buried nearer the surface of the soil experience more extreme temperatures and more temperature variation (Snell-Rood, Burger, Hutton, & Moczek, 2016), and the dung beetle gut microbiome can increase fitness under thermal stress (Schwab et al., 2016). Thus, if *P. vindex* buries its brood balls at shallower depths in sympatry than in allopatry, members of its gut microbiome may be locally adapted to more extreme temperatures in sympatry than in allopatry.

An alternative means of niche partitioning that could alter the gut microbiome is that *P. vindex*, an edaphic generalist throughout most of its range, is displaced by *P. difformis*, a sand-specialist, on sand where they co-occur (Blume & Aga 1976; 1978). However, this possibility seems unlikely because we consistently caught both species in high abundance in the same traps in sandy sites (Table S1).

We cannot say with certainty why the interaction between species and range overlap was significant in our Jaccard analysis but not our weighted UniFrac analysis. However, a lack of phylogenetic signal does not preclude the importance of the change in the gut microbiome, because physiologically important traits of gut microbes can differ among bacterial species, or even from strain to strain (e.g. Arnold, Simpson, Roach, Kwintkiewicz, & Azcarate-Peril, 2018). Future research should investigate differences in brood ball burial depths where *P. vindex* and *P. difformis* occur alone and in sympatry, coupled with metagenomic and metatranscriptomic approaches to understand if differences in members of the gut microbiome correspond to differences in functionality.

### 4.3 Local and broad scale trends shape gut microbial communities

Both the weighted UniFrac and the quantitative Jaccard models highlighted that geographic distance among *Phanaeus* communities, a few negative dbMEMs, and cattle abundance drove patterns of gut microbial variation (Figs. 5 and 6). However, our first variation partitioning analyses and model selections suggested that precipitation and temperature were also important predictors for gut microbial variation (Table S2, Table S3), but that they were colinear with geographic variables such as latitude and longitude (Fig. S7). Thus, we captured most of the contribution of longitude by including precipitation in our UniFrac model, whereas the effects of temperature variation were likely accounted for by including latitude in both the UniFrac and Jaccard models.

Lab studies on insects and other ectotherms have shown that temperature induces changes in the composition or function of the gut microbiome. As examples, insects under heat stress tend to have increases in the relative abundance of Proteobacteria in their guts (Sepulveda & Moeller, 2020) and the richness of salamander gut bacteria is substantially decreased at high experimental temperatures (Fontaine, Novarro, & Kohl, 2018). Elevated temperatures can also cause a loss of beneficial endosymbionts in aphids (Russell & Moran, 2006) and stinkbugs (Kikuchi et al., 2016). Research on temperature stress is essential for understanding the function of the gut microbiome and its potential response to climate change. However, we have a limited understanding of how temperature may interact with other selection pressures and host genetics across a species’ range to influence the gut microbiome. Other than two studies on the gut microbiota of the common fruit fly *Drosophila melanogaster* (Walters et al., 2020; Wang, Kapun, Waidele, Kuenzel, Bergland, & Staubach, 2020), ours is the only study to our knowledge that examines the gut microbiome of an insect across a temperature gradient representing much of its geographic range. We know of no other studies that examine how precipitation variation correlates with the gut microbiome. However, because the dung beetle gut microbiome leads to greater fitness outcomes under desiccation and temperature stressors (Schwab et al., 2016), it follows that selection would favor different gut microbial members under different precipitation and temperature regimes. Alternatively, changes in the gut microbiome may not reflect selection pressures on the host, but instead be the result of selection acting on free-living, environmental microbes encountered in the diet (Rosa, Minard, Lindholm, & Saastamoinen, 2019).

The significance of negative dbMEMs and cattle abundance in our models suggests that local factors shape the gut microbiome. In our system, negative spatial autocorrelation (i.e. significant negative dbMEMs) meant that dissimilar gut microbiome samples tended to cluster close to one another in space. Negative spatial autocorrelation is often the signature of biotic interactions (Borcard et al., 2018), and we found that among sympatric samples, the gut microbiome differs based on *Phanaeus* species identity (Fig. S9). Thus, this trend was likely due to structuring within sympatric sampling locations and among allopatric and sympatric *P. vindex* populations that were in close proximity to one another.

In contrast to negative dbMEMs, not a single positive dbMEM was significant, indicating that we did not detect positive spatial autocorrelation in our data. There are a few possible reasons for this. First, the differences in gut microbial communities with decreased distance among sites may be caused by temperature, precipitation, or other environmental variables that scale linearly with distance. In addition, the amount of cattle at a sampling site, which varied randomly across the landscape, was a strong predictor of gut microbiome function, suggesting factors related to diet are impacting the gut microbiome. As an example, the gut microbiome of beetles in areas with abundant cattle may be a result of priority effects, where the first microbes that beetles encounter in their diet (i.e. a brood ball made from cattle dung) are able to colonize the gut first with limited competition from other microbes, as has been found in the gut microbiomes of honeybees (Ellegaard & Engel, 2019). Another possibility is that microbes specialized to facilitate the digestion of cattle dung are selected for in the dung beetle gut, and they may even be passed on from one generation to the next. An exciting area for future research might be transgenerational studies to tease apart the degrees of vertical versus horizontal transmission characterizing ASVs associated with diet.

### 4.4 Important taxa of the gut microbiome

Our study confirms that abundant microbes in the gut of *Phanaeus* dung beetles are similar to those found in other genera of dung beetles, suggesting possibly beneficial roles of these microbes. Some of the most abundant bacterial families identified in *Phanaeus*, including Enterobacteriaceae, Comamonadaceae, Moraxellaceae, and Planococcaceae (Fig. 3), are also among the most prevalent in *Onthophagus taurus* (Estes et al., 2013; Hammer et al., 2016), *Aphodius fossor* (Hammer et al., 2016), *Euoniticellus intermedius*, and *E. triangulates* dung beetles (Shukla et al., 2016). Taxa present within these families likely aid dung beetles in digesting nutrient-poor dung. For example, some members of Enterobacteriaceae perform cellulose digestion and nitrogen fixation in fruit flies (Behar, Yuval, & Jurkevitch, 2005), while others assist bark beetles in uric acid recycling (Morales-Jiménez et al., 2013). *Dysgonomonas* has been found on multiple continents in several species of dung beetles in the genera *Onthophagus* and *Euoniticellus* (Parker, Newton, & Moczek, 2020), and members of the genus carry nitrogen fixation genes (Inoue et al., 2015) while others break down lignocellulose in the guts of termites (Sun, Yang, Zhang, Shen, & Ni, 2015). *Acinetobacter* is a widespread genus of aerobic bacteria that has been isolated from cattle rumen (Chang, Rhee, Jeong, Kim, & Kim, 2015) and insects such as red flour beetles (Wang, Xin, Shi, & Zhang, 2020). *Acinetobacter* may aid *Phanaeus* in breaking down difficult components of dung, as previous studies have shown members of the genus to digest recalcitrant polysterenes (Wang, Xin, Shi, & Zhang, 2020), lipids, and esters (Kok, Christoffels, Vosman, & Hellingwerf, 1993). Together our results and those from previous studies suggest that taxa in the genera *Acintobacter* and *Dysgonomnas* are likely vertically-transmitted, core members of the dung beetle microbiome.

### 4.5 Conclusion

This study offers a look into how biotic interactions and climatic factors across a species range interact to shape the gut microbiome. We found that the gut microbiomes of *Phanaeus vindex* and *P. difformis* vary across the species ranges in patterns predicted by cattle abundance in the local area, climatic factors such as precipitation and temperature, and biotic interactions. Range overlap and species identity were surprisingly poor predictors of variation among *Phanaeus* beetles in our full, pooled dataset. However, the gut microbiomes of *P. vindex* and *P. difformis* consistently exhibited turnover from allopatric populations to sympatric populations, and sympatric populations of *Phanaeus* harbored species-specific gut microbiomes. Furthermore, it appears that the gut microbiome of *P. vindex* may shift more between allopatric and sympatric populations than that of *P. difformis*, possibly indicating that the presence of *P. difformis* is driving character displacement of the gut microbiome of *P. vindex* where the species co-occur. However, more research is required to assess any physiological changes that may accompany the sympatric shifts in the gut microbiome and their effects on fitness. Our work emphasizes the roles of climatic variation, distance, and biotic interactions in shaping the gut microbiome and underscores the need to consider variation in the gut microbiome among populations in future work.

## ACKNOWLEDGEMENTS

We are grateful to Jim Fordyce for helpful discussions on statistical analyses and to Veronica Brown for advice on molecular techniques. We thank Todd Pierson, Sarah Jayne Brawner, and Brianna Jacobs for their assistance in the lab and for sourcing environmental data. We are grateful to Camilla Winfrey and Dane Winfrey for help with fieldwork. For access to trapping sites, we thank the University of Oklahoma Biological Station, the Oklahoma Department of Wildlife Conservation, the Texas Parks and Wildlife Department, the Konza Prairie Biological Station, the Fort Worth Nature Center and Refuge, and many private ranchers in Oklahoma and Texas. Funding was generously provided by the University of Tennessee, Knoxville (UTK), Sigma Xi, and the Coleopterists Society. C.C.W. acknowledges fellowship support from the National Science Foundation (Graduate Research Fellowship Program, grant number DGE 1650115) and from the Program for Excellence & Equity in Research, a UTK fellowship sponsored by the National Institutes of Health (IMSD, grant #R25GM086761). K.S.S. was funded by the US National Science Foundation (IOS-1930829).

## DATA ACCESSIBILITY

Raw 16S sequence data are available at the NCBI Sequence Read Archive (BioProject: SUB9081005). Code used for data processing and analysis will be made available upon peer-reviewed publication.

## AUTHOR CONTRIBUTIONS

C.C.W. and K.S.S. designed the study, based on an original idea conceived by K.S.S. C.C.W. collected samples, performed lab work, and analyzed the data. C.C.W. lead the writing of the manuscript, with substantial editing and revisions made by K.S.S.

---

**Figure S4.**
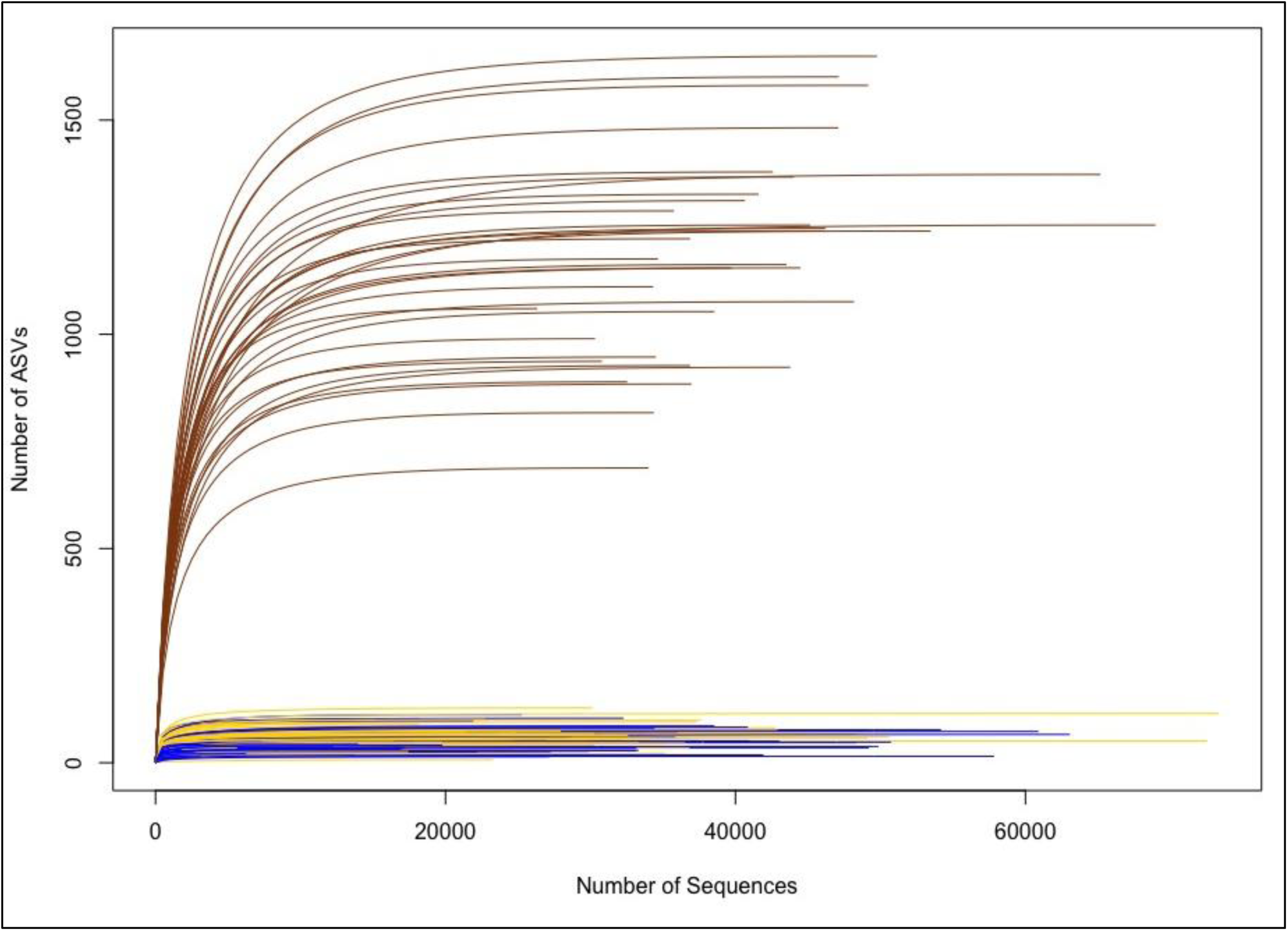
Rarefaction curves indicate that rarefaction depths chosen were adequate to capture ASV diversity in our samples. Brown lines represent soil samples, whereas blue and gold lines are *P. vindex* and *P. difformis* samples, respectively. Soil samples were rarefied at 26,336 reads and gut samples were rarefied at 3,500 reads because soils were far more diverse than gut samples and overall had better sampling coverage.

**Figure S7:**
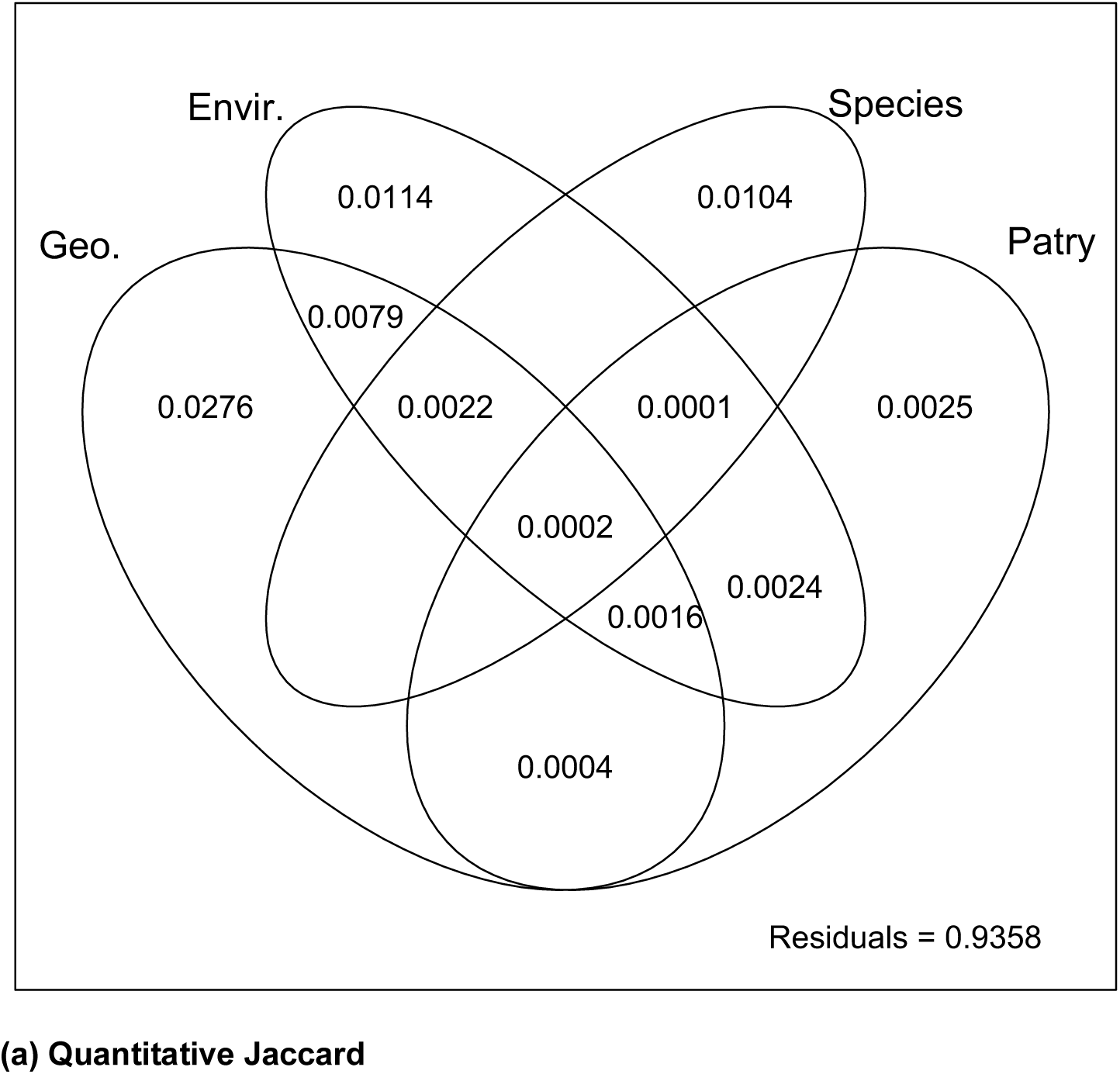
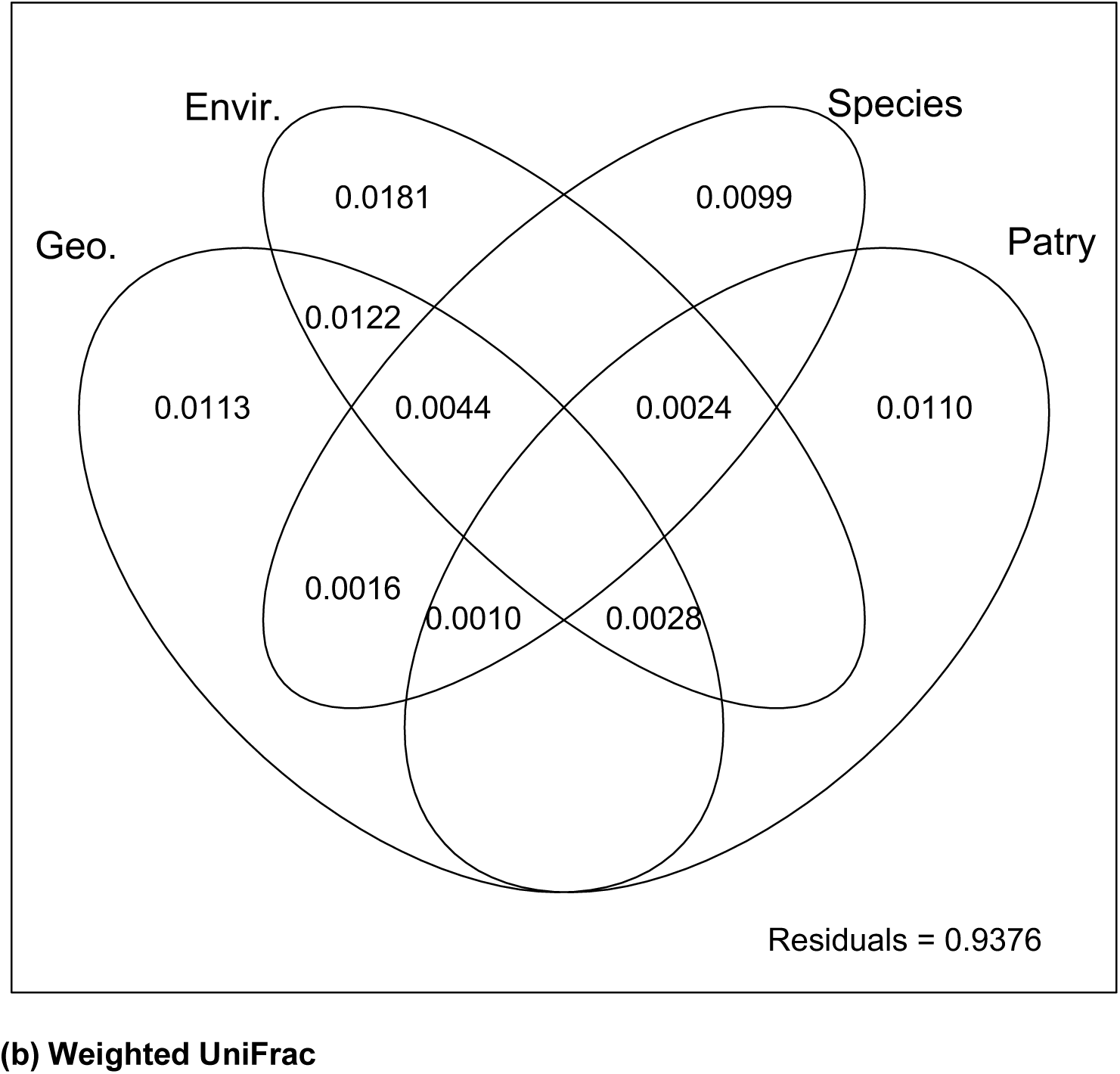
Variation partitioning of *Phanaeus vindex* and *P. difformis* gut microbiome data into various components. Response variables are (a) quantitative Jaccard dissimilarity and (b) weighted UniFrac distances. In both plots, Species refers to the amount of variation explained by *Phanaeus* species (*P. vindex* or *P.* difformis) and Patry refers to range overlap (sympatry or allopatry of *Phanaeus* species). In (a), geographic variables (Geo.) include X,Y coordinates (longitude and latitude) and six negative distance-based Moran’s eigenvector mapping variables (MEMs 2,5,8,10,11, and 12); environmental variables (Envir.) include mean temperature, mean precipitation, the mean Shannon index of soil samples taken from the sampling area, and the amount of cattle present. In (b), Geo includes X,Y coordinates and one negative MEM (MEM 6); and Envir. variables include mean temperature, mean precipitation, and abundance of cattle present. Numbers within the Venn diagram represent the adjusted *R^2^* of each fraction. Blank fractions have very small negative adjusted *R^2^* values and are not shown.

**Figure S8:**
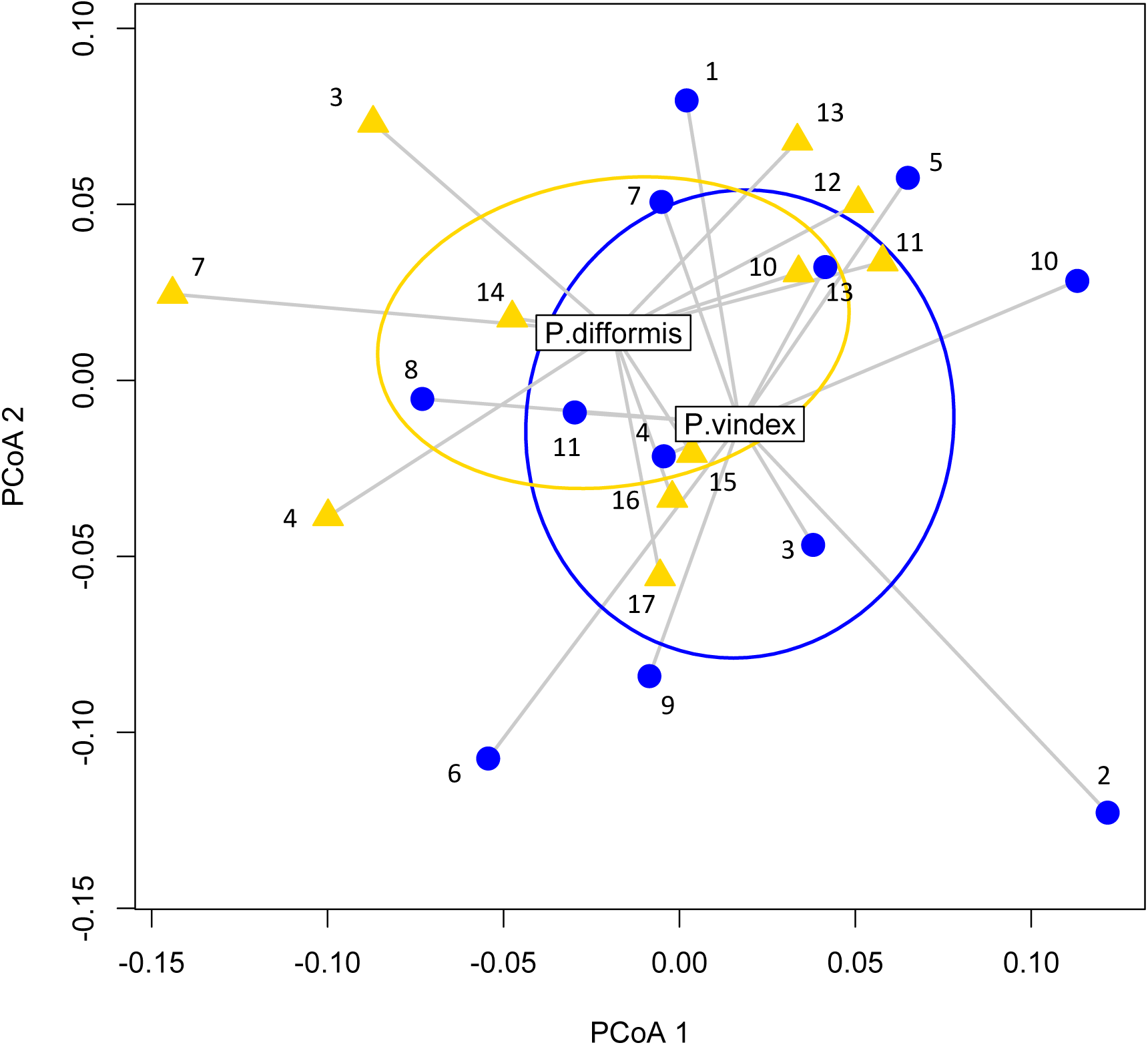
Beta diversity of *Phanaeus vindex* and *P. difformis* represented as multivariate dispersions based on weighted UniFrac distances. Populations of *P. vindex* are represented by blue dots; gold triangles indicate populations of *P. difformis*. Numbers represent different populations by location (see Table S1 for metadata associated with each population). The words “*P. vindex*” and “*P. difformis*” are positioned in the species’ respective centroids in multivariate space. Ellipses represent one standard deviation away from the centroid of *P. vindex* and *P. difformis*.

**Figure S9:**
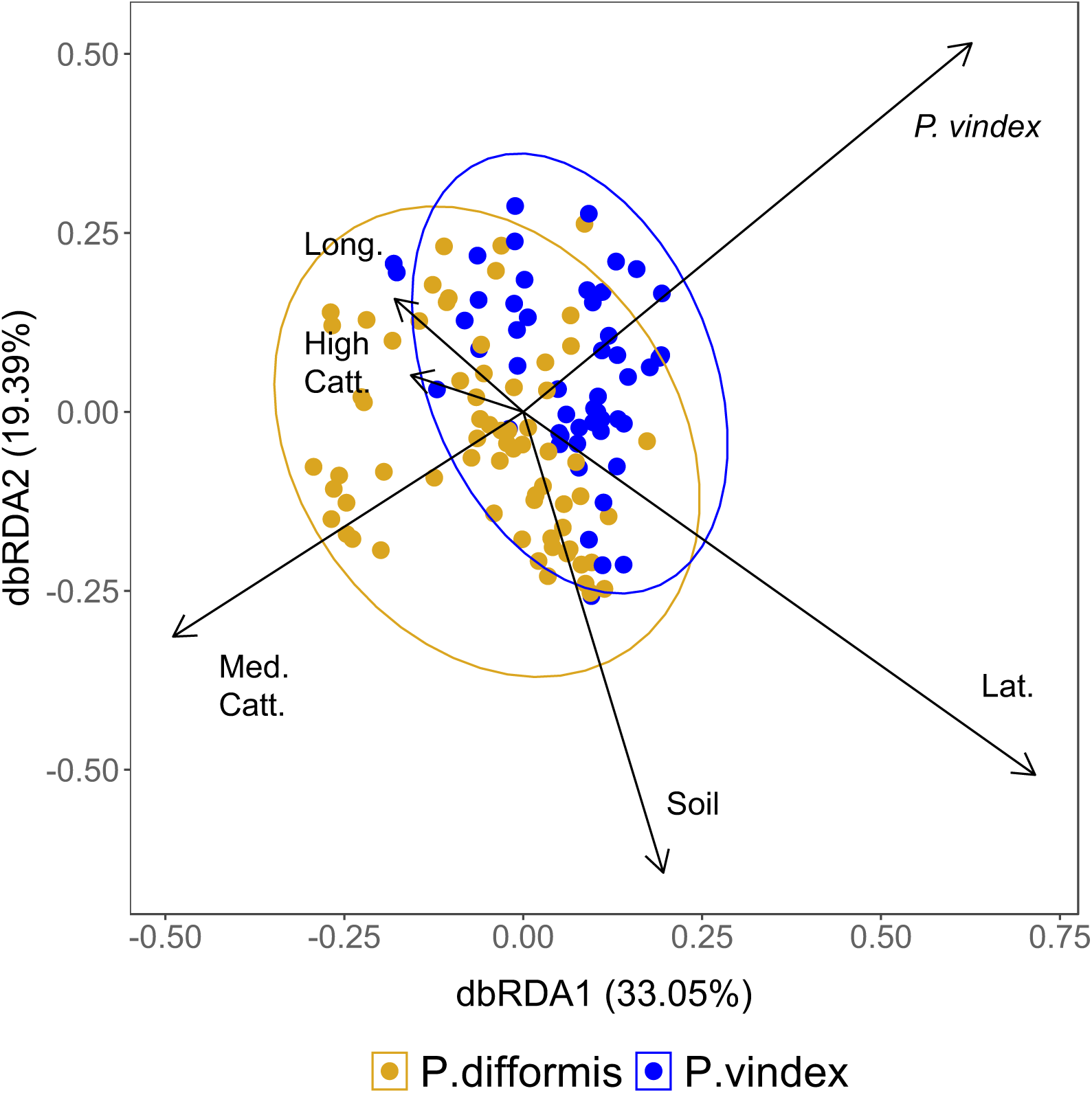
Plot of distance-based redundancy analysis (db-RDA) based on the quantitative Jaccard dissimilarity among samples of *Phanaeus vindex* and *P. difformis* in sympatry. Points on the plot indicate individual *P. difformis* and *P. vindex* beetles. Ellipses represent 95% confidence intervals around centroids of *P. difformis* in allopatry and in sympatry with *P. vindex*. All constrained axes together explain 11.24% of the total variation in multivariate space. Arrows indicate the strength of correlation of variables with dbRDA axes 1 and 2. Numerical variables constraining the ordination latitude (Lat), longitude (Long.), and the mean Shannon index of soil samples taken from the sampling areas (Soil). Constraining factor variables include cattle presence in the sampling area (low, medium, or high), and range overlap (i.e. sympatry or allopatry). Factor variables are automatically dummy coded; thus, the factor level coded as the intercept does not have a corresponding axis. All variables shown were significant following ANOVAs (99,999 permutations).

**Table S1.**
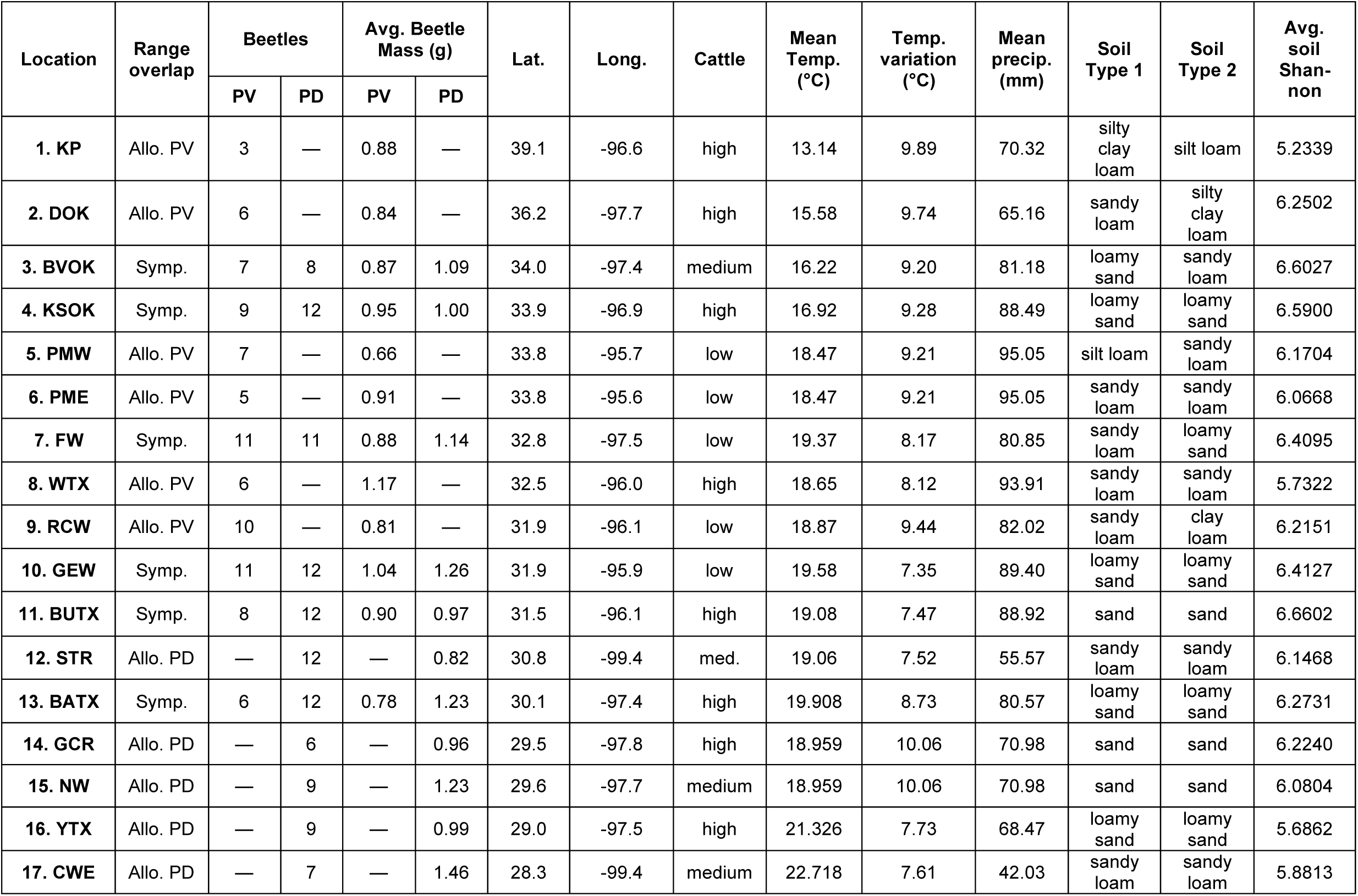
Metadata associated with each sampling location. PV and PD refer to *Phanaeus vindex* and *P. difformis*, respectively. Allo. and symp. stand for allopatry and sympatry, respectively. Latitude and longitude reported here are to the nearest tenth of a degree to respect the privacy of landowners. Mean beetle mass refers to that of the beetle samples retained after rarefying. Mean temperature, temperature variation, and mean precipitation are based on mean monthly data over ten years. We used the United States Department of Agriculture’s (USDA) Web Soil Survey to assign the soil at each successful trap to one of the twelve major soil textural classes based on proportions of silt, sand, and clay as defined by the USDA (Soil Science Division Staff, 2017). Soil types reported here are the two most common soil types at each location (some locations had up to 3 soil types).

| Location | Range overlap | Beetles |  | Avg. Beetle Mass (g) |  | Lat. | Long. | Cattle | Mean Temp. (°C) | Temp. variation (°C) | Mean precip. (mm) | Soil Type 1 | Soil Type 2 | Avg. soil Shan-non |
| --- | --- | --- | --- | --- | --- | --- | --- | --- | --- | --- | --- | --- | --- | --- |
|  |  | PV | PD | PV | PD |  |  |  |  |  |  |  |  |  |
| 1. KP | Allo. PV | 3 | — | 0.88 | — | 39.1 | -96.6 | high | 13.14 | 9.89 | 70.32 | silty clay loam | silt loam | 5.2339 |
| 2. DOK | Allo. PV | 6 | — | 0.84 | — | 36.2 | -97.7 | high | 15.58 | 9.74 | 65.16 | sandy loam | silty clay loam | 6.2502 |
| 3. BVOK | Symp. | 7 | 8 | 0.87 | 1.09 | 34.0 | -97.4 | medium | 16.22 | 9.20 | 81.18 | loamy sand | sandy loam | 6.6027 |
| 4. KSOK | Symp. | 9 | 12 | 0.95 | 1.00 | 33.9 | -96.9 | high | 16.92 | 9.28 | 88.49 | loamy sand | loamy sand | 6.5900 |
| 5. PMW | Allo. PV | 7 | — | 0.66 | — | 33.8 | -95.7 | low | 18.47 | 9.21 | 95.05 | silt loam | sandy loam | 6.1704 |
| 6. PME | Allo. PV | 5 | — | 0.91 | — | 33.8 | -95.6 | low | 18.47 | 9.21 | 95.05 | sandy loam | sandy loam | 6.0668 |
| 7. FW | Symp. | 11 | 11 | 0.88 | 1.14 | 32.8 | -97.5 | low | 19.37 | 8.17 | 80.85 | sandy loam | loamy sand | 6.4095 |
| 8. WTX | Allo. PV | 6 | — | 1.17 | — | 32.5 | -96.0 | high | 18.65 | 8.12 | 93.91 | sandy loam | sandy loam | 5.7322 |
| 9. RCW | Allo. PV | 10 | — | 0.81 | — | 31.9 | -96.1 | low | 18.87 | 9.44 | 82.02 | sandy loam | clay loam | 6.2151 |
| 10. GEW | Symp. | 11 | 12 | 1.04 | 1.26 | 31.9 | -95.9 | low | 19.58 | 7.35 | 89.40 | loamy sand | loamy sand | 6.4127 |
| 11. BUTX | Symp. | 8 | 12 | 0.90 | 0.97 | 31.5 | -96.1 | high | 19.08 | 7.47 | 88.92 | sand | sand | 6.6602 |
| 12. STR | Allo. PD | — | 12 | — | 0.82 | 30.8 | -99.4 | med. | 19.06 | 7.52 | 55.57 | sandy loam | sandy loam | 6.1468 |
| 13. BATX | Symp. | 6 | 12 | 0.78 | 1.23 | 30.1 | -97.4 | high | 19.908 | 8.73 | 80.57 | loamy sand | loamy sand | 6.2731 |
| 14. GCR | Allo. PD | — | 6 | — | 0.96 | 29.5 | -97.8 | high | 18.959 | 10.06 | 70.98 | sand | sand | 6.2240 |
| 15. NW | Allo. PD | — | 9 | — | 1.23 | 29.6 | -97.7 | medium | 18.959 | 10.06 | 70.98 | sand | sand | 6.0804 |
| 16. YTX | Allo. PD | — | 9 | — | 0.99 | 29.0 | -97.5 | high | 21.326 | 7.73 | 68.47 | loamy sand | loamy sand | 5.6862 |
| 17. CWE | Allo. PD | — | 7 | — | 1.46 | 28.3 | -99.4 | medium | 22.718 | 7.61 | 42.03 | sandy loam | sandy loam | 5.8813 |

**Table S2.** Results of forward model selection based on adjusted *R^2^* and *P*-values, performed prior to variation partitioning. Jaccard dissimilarities and weighted UniFrac dissimilarities were used as response variables. (a) Model selection on negative distance-based Moran’s eigenvector map (dbMEM) variables. Because a linear trend was detected, detrended dissimilarity matrices were used as the response variables. (b) Model selection on environmental variables.

**(a) Model selection on negative distance-based Moran's eigenvector map variables**
| Dissimilarity matrix | MEM: | After double-stopping forward model selection |  | After Sidák correction |
| --- | --- | --- | --- | --- |
| | | Adjusted $R^2$ | $P$-value | Adjusted $p$-value |
| Jaccard | MEM1 | 0.036909 | < 0.05 * | 0.19132 |
|  | MEM2 | 0.016477 | < 0.01 ** | < 0.05 * |
|  | MEM5 | 0.024535 | < 0.01 ** | < 0.05 * |
|  | MEM6 | 0.03904 | < 0.05 * | 0.29557 |
|  | MEM7 | 0.034445 | < 0.01 ** | 0.09164 |
|  | MEM8 | 0.028196 | < 0.01 ** | < 0.05 * |
|  | MEM9 | 0.031439 | < 0.01 ** | 0.06198 |
|  | MEM10 | 0.006983 | < 0.0001 *** | < 0.001 *** |
|  | MEM11 | 0.012124 | < 0.001 *** | < 0.01 ** |
|  | MEM12 | 0.020868 | < 0.01 ** | < 0.05 * |
| Weighted UniFrac | MEM6 | 0.010804 | < 0.01 ** | < 0.05 * |
|  | MEM7 | 0.019963 | < 0.05 * | 0.06797 |
|  | MEM8 | 0.033577 | < 0.05 * | 0.11256 |
|  | MEM11 | 0.027603 | < 0.05 * | 0.19922 |

**(b). Model selection on environmental variables**
| Dissimilarity matrix | Environmental variable | After double-stopping forward model selection |  |
| --- | --- | --- | --- |
| | | Adjusted $R^2$ | $P$-value |
| **Jaccard** | Cattle presence | 0.010721 | 0.0001 *** |
|  | Mean soil Shannon entropy | 0.015704 | < 0.001 *** |
|  | Mean monthly temperature | 0.020614 | < 0.001 *** |
|  | Mean monthly precipitation | 0.025039 | < 0.01 ** |
|  | All variables | 0.027232 | N/A |
| **weighted UniFrac** | Cattle presence | 0.020423 | < 0.05 * |
|  | Mean monthly temperature | 0.012315 | < 0.01 ** |
|  | Mean monthly precipitation | 0.033710 | < 0.01 ** |
|  | All variables | 0.038557 | N/A |

**Table S3.** Results of forward model selection based on adjusted *R^2^* and *P*-value, performed prior to implementing distance-based redundancy analyses (db-RDAs) and additional variation partitioning. Quantitative Jaccard and weighted UniFrac distances were used as response variables. All variables except precipitation in the Jaccard model were included in final db-RDAs and second round of variation partitioning analyses.

|  |  | After double-stopping forward model selection |  |
| --- | --- | --- | --- |
| Dissimilarity matrix | Predictor | Adjusted $R^2$ | $P$-value |
| Jaccard | X coordinates (longitude) | 0.062416 | < 0.01 ** |
|  | Y coordinates (latitude) | 0.037077 | < 0.001 *** |
|  | Negative dbMEM 2 | 0.053218 | < 0.05 * |
|  | Negative dbMEM 5 | 0.050680 | < 0.01 ** |
|  | Negative dbMEM 8 | 0.057153 | < 0.01 ** |
|  | Negative dbMEM 10 | 0.046759 | < 0.01 ** |
|  | Negative dbMEM 11 | 0.032073 | < 0.001 *** |
|  | Ranked cattle abundance | 0.010721 | < 0.001 *** |
|  | Mean precipitation | 0.042212 | < 0.001 *** |
|  | Mean Shannon entropy of soil | 0.058946 | < 0.05 * |
|  | *Phanaeus* species | 0.019942 | < 0.001 *** |
|  | Range overlap ("patry") | 0.026400 | < 0.001 *** |
|  | All variables | 0.064215 | N/A |
| Weighted UniFrac | Y coordinates (latitude) | 0.016370 | < 0.001 *** |
|  | Negative dbMEM 6 | 0.060728 | < 0.05 * |
|  | Ranked cattle abundance | 0.037549 | < 0.05 * |
|  | Mean precipitation | 0.053021 | < 0.05 * |
|  | *Phanaeus* species | 0.044704 | < 0.05 * |
|  | Range overlap ("patry") | 0.028341 | < 0.01 ** |
|  | All variables | 0.064637 | N/A |

**Table S5.**
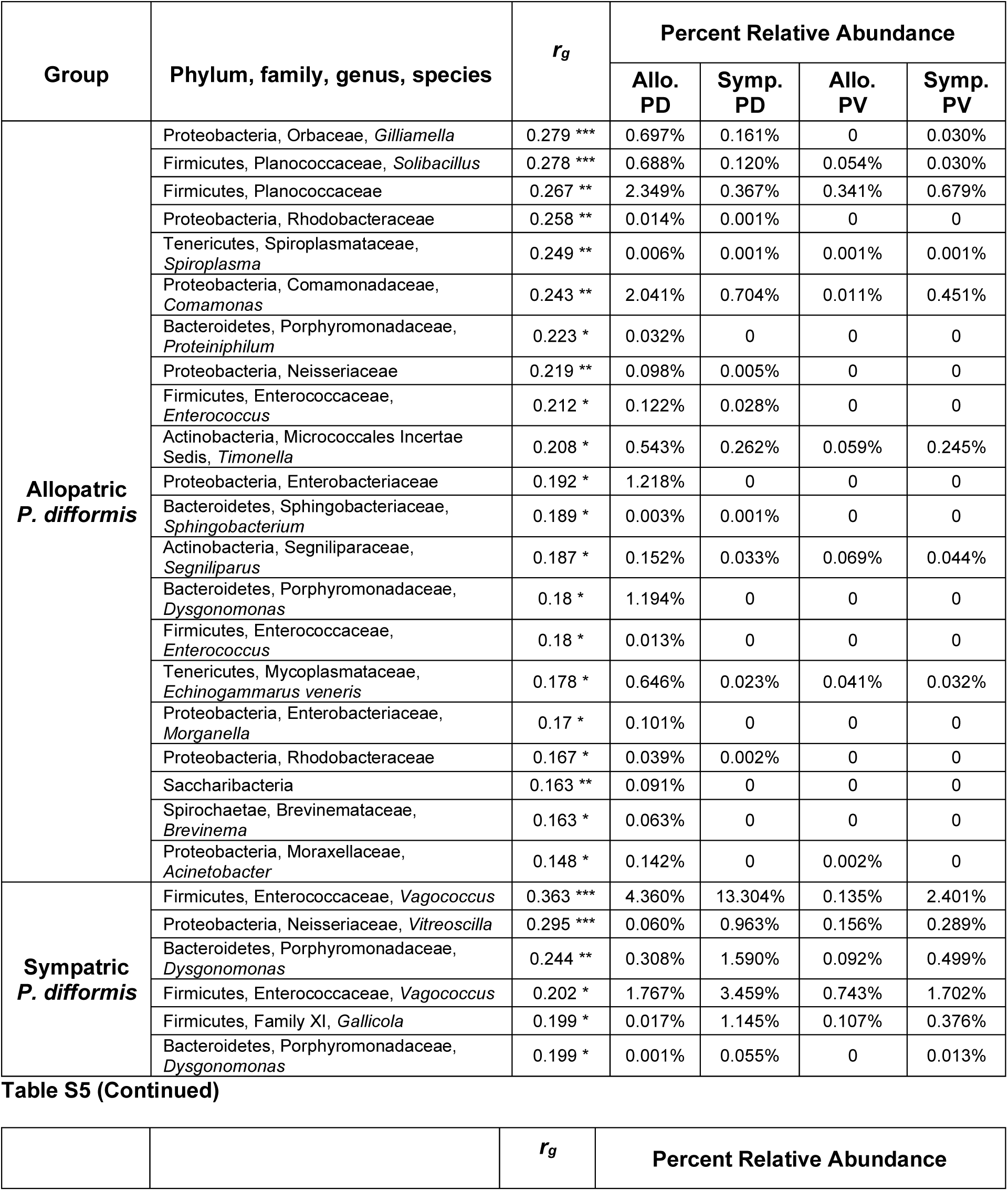
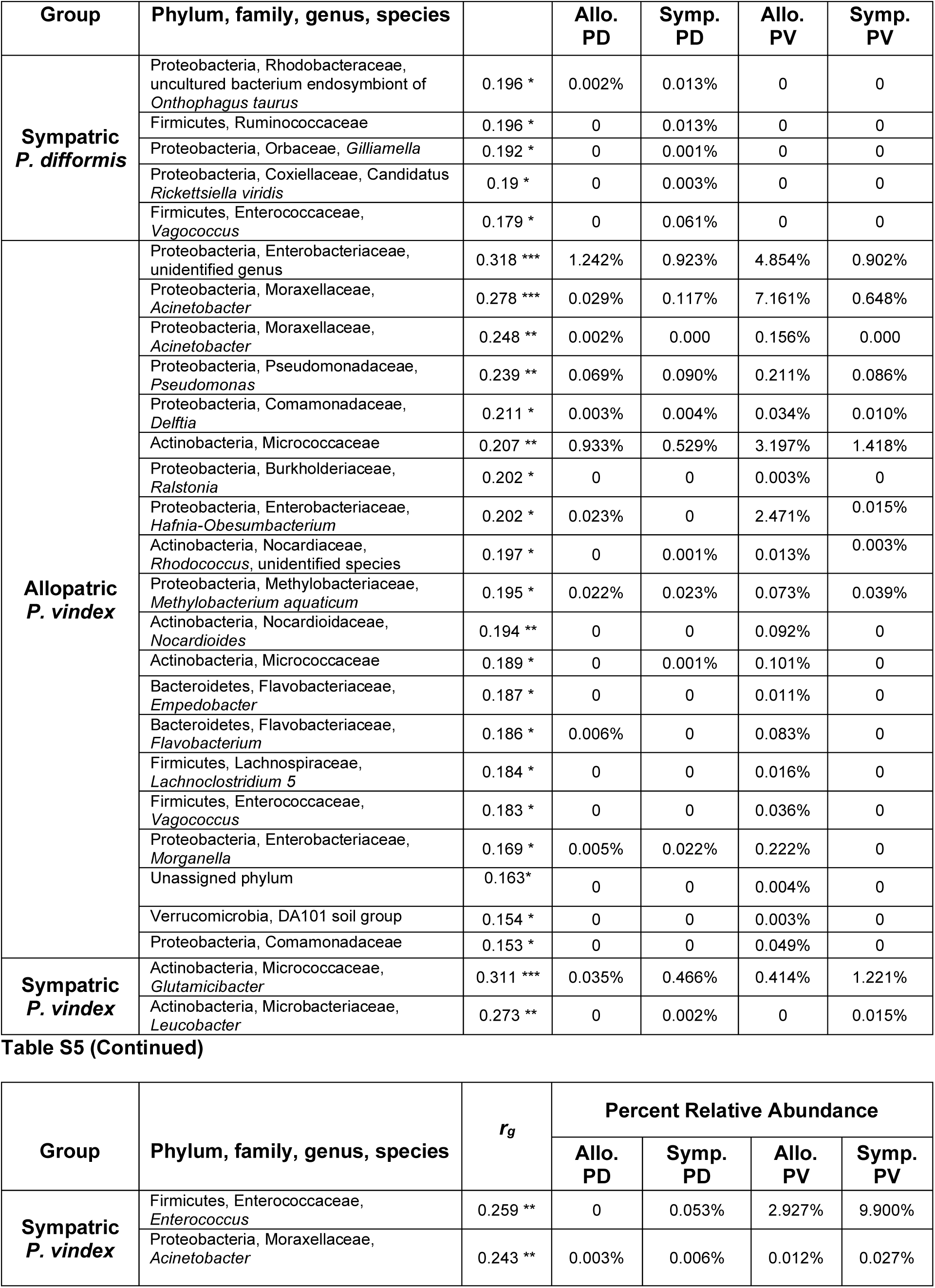
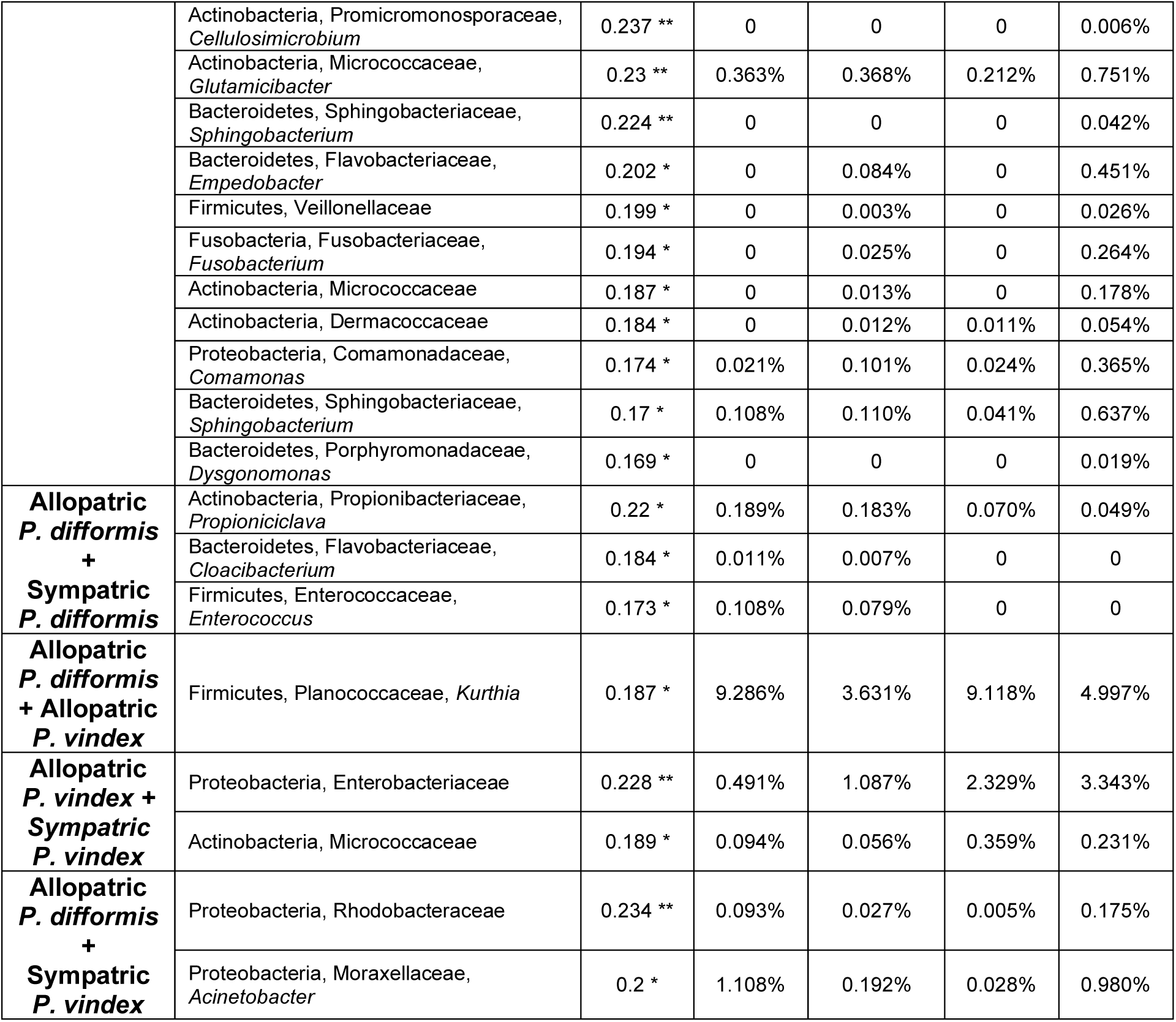
Indicator species analyses for different groupings of *Phanaeus difformis* and *P. vindex* gut microbiome samples in allopatry and in sympatry. Taxonomic information is presented as phylum, family, genus, species, as available in the SILVA 128 99% OTUs reference database. Asterisks next to *r_g_* values indicate significance levels; * *p* < 0.05, ** *p* < 0.01, and *** *p* < 0.001. PD and PV stand for *P. difformis* and *P. vindex*, respectively. No ASVs characterizing the combination of sympatric *P. vindex* and sympatric *P. difformis* gut microbiome samples were identified.

**Table S6.**
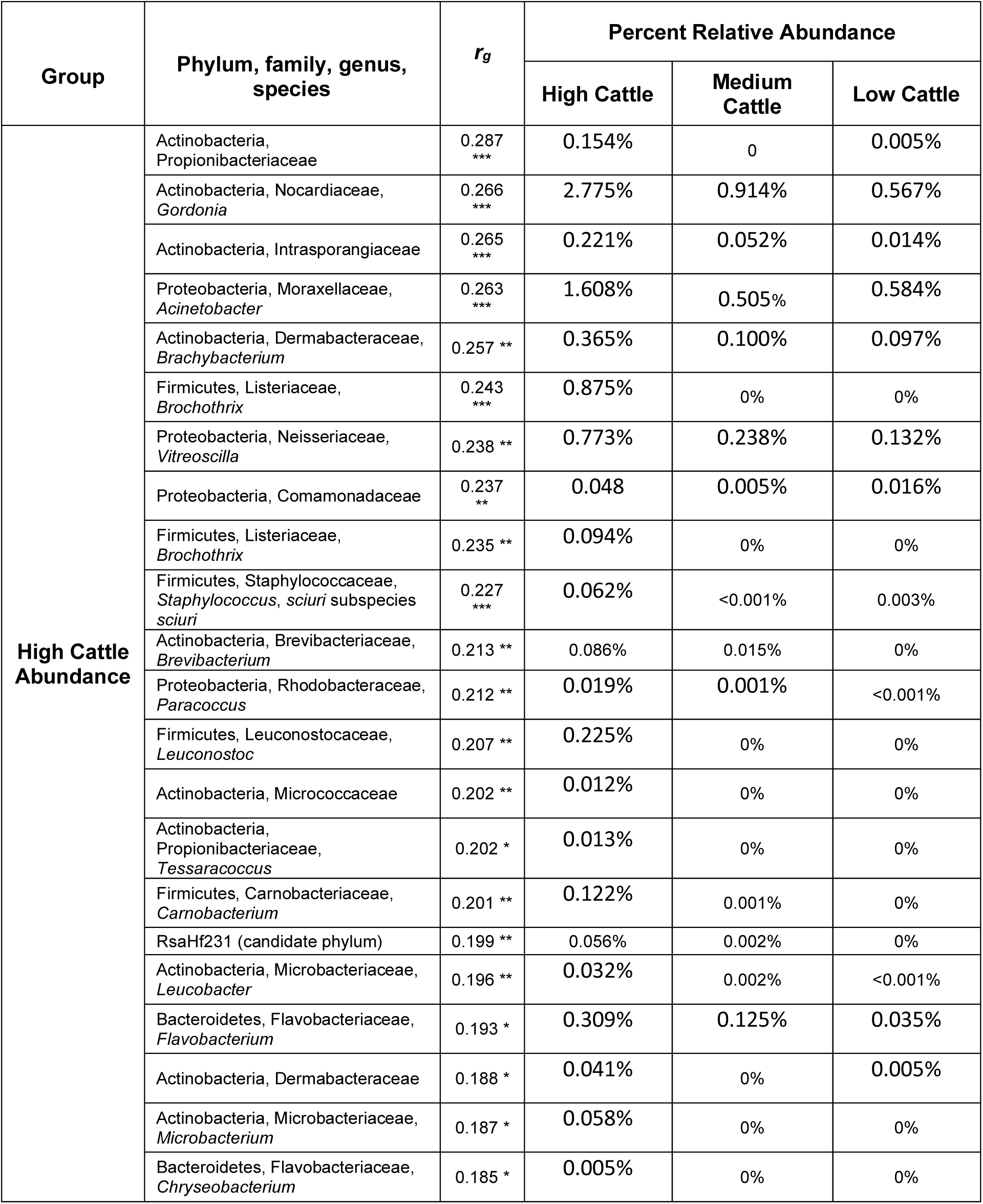
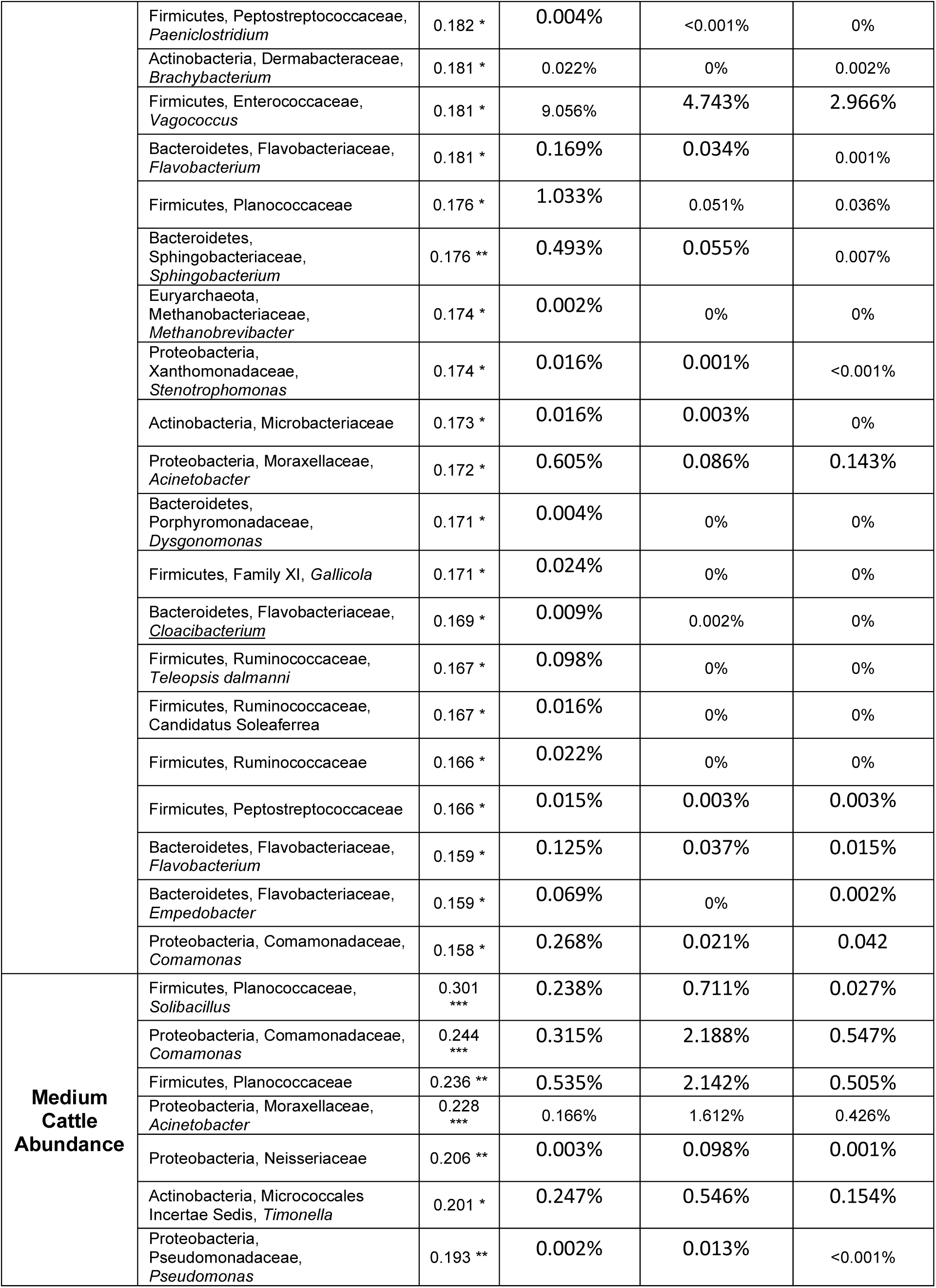
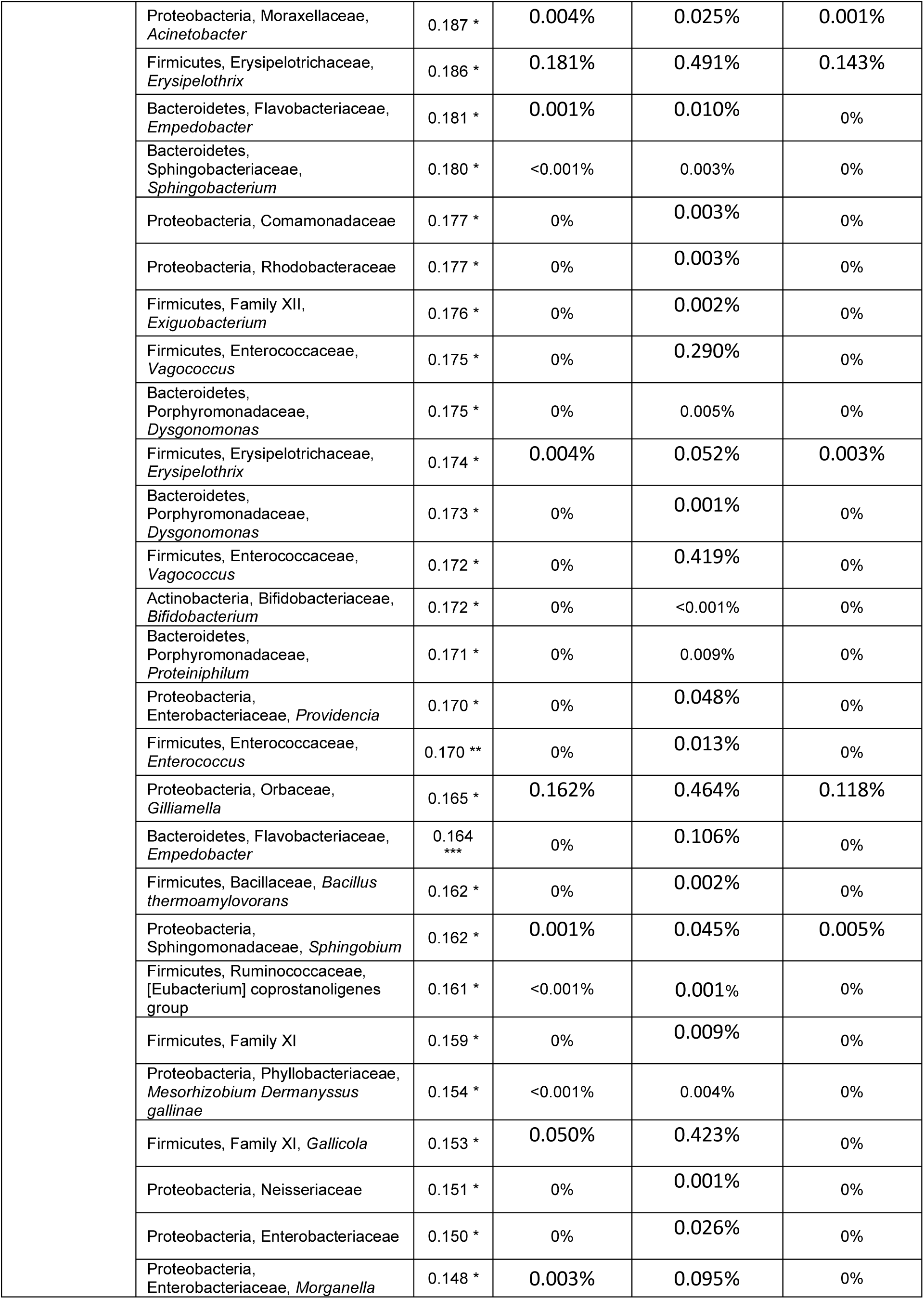
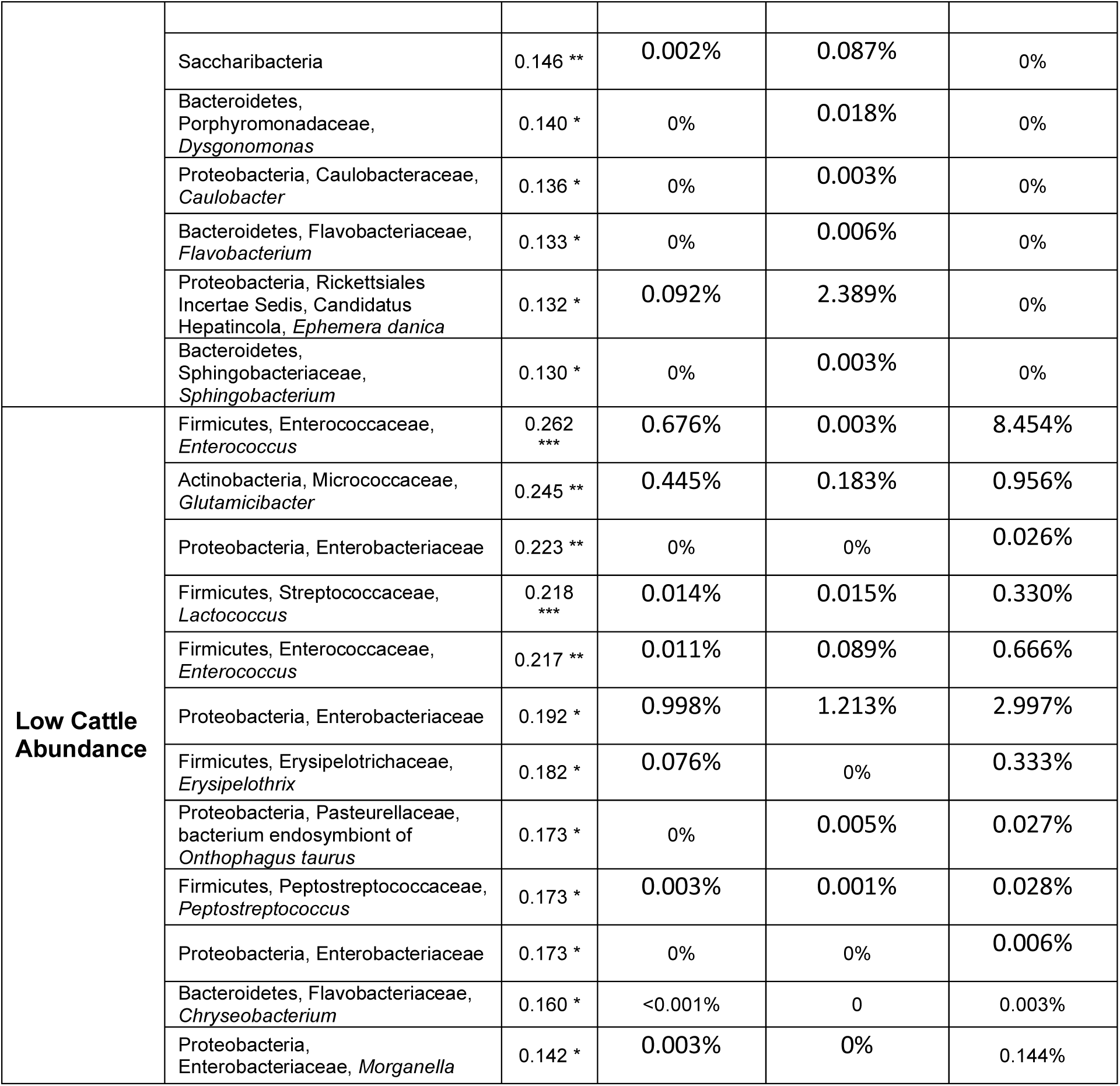
Indicator species analyses for the gut microbial communities of *Phanaeus vindex* and *P. difformis* collected in areas with high, medium, or low cattle abundance. Taxonomic information is presented as phylum, family, genus, species, as available in the SILVA 128 99% OTUs reference database. Asterisks next to *r_g_* values indicate significance levels; * *p* < 0.05, ** *p* < 0.01, and *** *p* < 0.001. PD and PV stand for *P. difformis* and *P. vindex*, respectively. No ASVs characterizing combinations of different cattle abundances were identified.

## REFERENCES

1. Anderson, M. J., Ellingsen, K. E., & McArdle, B. H. (2006). Multivariate dispersion as a measure of beta diversity. Ecology Letters, 9(6), 683–693. doi: 10.1111/j.1461-0248.2006.00926.x

2. Apprill, A., McNally, S., Parsons, R., & Weber, L. (2015). Minor revision to V4 region SSU rRNA 806R gene primer greatly increases detection of SAR11 bacterioplankton. Aquatic Microbial Ecology, 75(2), 129–137. doi: https://doi.org/10.3354/ame01753

3. Arnold, J. W., Simpson, J. B., Roach, J., Kwintkiewicz, J., & Azcarate-Peril, M. A. (2018). Intra-species genomic and physiological variability impact stress resistance in strains of probiotic potential. Frontiers in Microbiology, 9, 242. doi: 10.3389/fmicb.2018.00242

4. Behar, A., Yuval, B., & Jurkevitch, E. (2005). Enterobacteria-mediated nitrogen fixation in natural populations of the fruit fly *Ceratitis capitata*. Molecular Ecology, 14(9), 2637–2643. doi: 10.1111/j.1365-294X.2005.02615.x

5. Benson, A. K., Kelly, S. A., Legge, R., Ma, F., Low, S. J., Kim, J., … & Kachman, S. D. (2010). Individuality in gut microbiota composition is a complex polygenic trait shaped by multiple environmental and host genetic factors. Proceedings of the National Academy of Sciences, 107(44), 18933–18938. doi: 10.1073/pnas.1007028107

6. Blanchet, F. G., Legendre, P., & Borcard, D. (2008). Forward selection of explanatory variables. Ecology, 89(9), 2623–2632. doi: 10.1890/07-0986.1

7. Blume, R. R., & Aga, A. (1976). *Phanaeus difformis* LeConte (Coleoptera: Scarabaeidae): Clarification of published descriptions, notes on biology, and distribution in Texas. *The Coleopterists’* Bulletin, 199–205.

8. Blume, R. R., & Aga, A. (1978). Observations on ecological and phylogenetic relationships of *Phanaeus difformis* LeConte and *Phanaeus vindex* MacLeay (Coleoptera: Scarabaeidae) in North America. Southwest Entomology, 3(2), 113–120.

9. Bolyen, E., Rideout, J. R., Dillon, M. R., Bokulich, N. A., Abnet, C. C., Al-Ghalith, G. A., … & Bai, Y. (2019). Reproducible, interactive, scalable and extensible microbiome data science using QIIME 2. Nature Biotechnology, 37(8), 852–857. doi: 10.1038/s41587-019-0209-9

10. Borcard, D., & Legendre, P. (2002). All-scale spatial analysis of ecological data by means of principal coordinates of neighbour matrices. Ecological Modelling, 153(1-2), 51–68. doi: 10.1016/S0304-3800(01)00501-4

11. Borcard, D., Gillet, F., & Legendre, P. (2018). Numerical Ecology with R (2nd ed.) New York, NY: Springer.

12. Borcard, D., Legendre, P., & Drapeau, P. (1992). Partialling out the spatial component of ecological variation. Ecology, 73(3), 1045–1055. doi: 10.2307/1940179

13. Borcard, D., Legendre, P., Avois-Jacquet, C., & Tuomisto, H. (2004). Dissecting the spatial structure of ecological data at multiple scales. Ecology, 85(7), 1826–1832. doi: 10.1890/03-3111

14. Bordenstein, S. R., & Theis, K. R. (2015). Host biology in light of the microbiome: ten principles of holobionts and hologenomes. PloS Biology, 13(8), e1002226. doi: 10.1371/journal.pbio.1002226

15. Bright, M., & Bulgheresi, S. (2010). A complex journey: transmission of microbial symbionts. Nature Reviews Microbiology, 8(3), 218. doi: 10.1038/nrmicro2262

16. Brown, W. L., & Wilson, E. O. (1956). Character displacement. Systematic Zoology, 5(2), 49–64. doi: 10.2307/2411924

17. Brucker, R. M., & Bordenstein, S. R. (2013). The hologenomic basis of speciation: gut bacteria cause hybrid lethality in the genus *Nasonia*. Science, 341(6146), 667–669. doi: 10.1126/science.1256708

18. Callahan, B. J., McMurdie, P. J., Rosen, M. J., Han, A. W., Johnson, A. J. A., & Holmes, S. P. (2016). DADA2: high-resolution sample inference from Illumina amplicon data. Nature Methods, 13(7), 581. doi: 10.1038/nmeth.3869

19. Caporaso, J. G., Lauber, C. L., Walters, W. A., Berg-Lyons, D., Lozupone, C. A., Turnbaugh, P. J., … & Knight, R. (2011). Global patterns of 16S rRNA diversity at a depth of millions of sequences per sample. Proceedings of the National Academy of Sciences, 108, 4516–4522. doi: 10.1073/pnas.1000080107

20. Carrier, T. J., & Reitzel, A. M. (2017). The hologenome across environments and the implications of a host-associated microbial repertoire. Frontiers in microbiology, 8, 802. doi.org/10.3389/fmicb.2017.00802

21. Chang, D. H., Rhee, M. S., Jeong, H., Kim, S., & Kim, B. C. (2015). Draft genome sequence of *Acinetobacter* sp. HR7, isolated from Hanwoo, Korean Native Cattle. Genome Announcements, 3(1). doi: 10.1128/genomeA.01358-14

22. Coon, K. L., Brown, M. R., & Strand, M. R. (2016). Mosquitoes host communities of bacteria that are essential for development but vary greatly between local habitats. Molecular Ecology, 25(22), 5806–5826. doi: 10.1111/mec.13877

23. Corbin, C., Heyworth, E. R., Ferrari, J., & Hurst, G. D. (2017). Heritable symbionts in a world of varying temperature. Heredity, 118(1), 10. doi: 10.1038/hdy.2016.71

24. De Cáceres, M. D., & Legendre, P. (2009). Associations between species and groups of sites: indices and statistical inference. Ecology, 90(12), 3566–3574. doi: 10.1890/08-1823.1

25. De Cáceres, M., Legendre, P., & Moretti, M. (2010). Improving indicator species analysis by combining groups of sites. Oikos, 119(10), 1674–1684. doi: 10.1111/j.1600-0706.2010.18334.x

26. Dickey, A. M. (2006). *Population genetics of* Phanaeus vindex *and* P. difformis *and congruence with morphology across a geographic zone of species overlap* (Unpublished MS thesis). The University of Texas at Arlington.

27. Dickey, J., Swenie, R., Turner, S., Winfrey, C., Yaffar, D., Padukone, A., … Kivlin, K. (2020). Do microorganisms obey macroecological rules?. Authorea. doi: 10.22541/au.159551320.05175629/v2

28. Dray, S., Bauman, D., Blanchet, G., Borcard, D., Clappe, S., Guenard, G., … & Wagner, H.H. (2020). adespatial: Multivariate Multiscale Spatial Analysis. R package version 0.3-8. Retrieved from https://CRAN.R-project.org/package=adespatial

29. Dray, S., Legendre, P., & Peres-Neto, P. R. (2006). Spatial modelling: a comprehensive framework for principal coordinate analysis neighbour matrices (PCNM). Ecological Modelling, 196(3-4), 483–493. doi: 10.1016/j.ecolmodel.2006.02.015

30. Dunbar, H. E., Wilson, A. C., Ferguson, N. R., & Moran, N. A. (2007). Aphid thermal tolerance is governed by a point mutation in bacterial symbionts. PLoS Biology, 5(5). doi: 10.1371/journal.pbio.0050096

31. Eddy, S. R. (2011). Accelerated profile HMM searches. PLoS Computational Biology, 7(10), e1002195. doi: 10.1371/journal.pcbi.1002195

32. Edmonds, W. D. (1994). Revision of *Phanaeus* Macleay, A New World Genus of Scarabaeinae Dung Beetles (Coleoptera: Scarabaeidae, Scarabaeinae). Natural History Museum of Los Angeles County Contributions in Science, 443, 1–105.

33. Ellegaard, K. M., & Engel, P. (2019). Genomic diversity landscape of the honey bee gut microbiota. Nature Communications, 10(1), 1–13. doi: 10.1038/s41467-019-08303-0

34. Engel, P., & Moran, N. A. (2013). The gut microbiota of insects–diversity in structure and function. FEMS Microbiology Reviews, 37(5), 699–735. doi: 10.1111/1574-6976.12025

35. Estes, A. M., Hearn, D. J., Snell-Rood, E. C., Feindler, M., Feeser, K., Abebe, T., … & Moczek, A. P. (2013). Brood ball-mediated transmission of microbiome members in the dung beetle, *Onthophagus taurus* (Coleoptera: Scarabaeidae). PLoS ONE, 8(11), e79061. doi: 10.1371/journal.pone.0079061

36. Fietz, K., Hintze, C. O. R., Skovrind, M., Nielsen, T. K., Limborg, M. T., Krag, M. A., … & Gilbert, M. T. P. (2018). Mind the gut: genomic insights to population divergence and gut microbial composition of two marine keystone species. Microbiome, 6(1), 1–16. doi: 10.1186/s40168-018-0467-7

37. Finn, R. D., Clements, J., & Eddy, S. R. (2011). HMMER web server: interactive sequence similarity searching. Nucleic Acids Research, 39(suppl_2), W29–W37. doi: 10.1093/nar/gkr367

38. Fontaine, S. S., Novarro, A. J., & Kohl, K. D. (2018). Environmental temperature alters the digestive performance and gut microbiota of a terrestrial amphibian. Journal of Experimental Biology, 221(20). doi: 10.1242/jeb.187559

39. Friedman, N., Shriker, E., Gold, B., Durman, T., Zarecki, R., Ruppin, E., & Mizrahi, I. (2017). Diet-induced changes of redox potential underlie compositional shifts in the rumen archaeal community. Environmental Microbiology, 19(1), 174–184. doi: 10.1111/1462-2920.13551

40. Hallali, E., Kokou, F., Chourasia, T. K., Nitzan, T., Con, P., Harpaz, S., … & Cnaani, A. (2018). Dietary salt levels affect digestibility, intestinal gene expression, and the microbiome, in Nile tilapia (Oreochromis niloticus). PloS ONE, 13(8), e0202351. doi: 10.1371/journal.pone.0202351

41. Hammer, T. J., Dickerson, J. C., & Fierer, N. (2015). Evidence-based recommendations on storing and handling specimens for analyses of insect microbiota. PeerJ, 3, e1190. doi: 10.7717/peerj.1190

42. Hammer, T. J., Fierer, N., Hardwick, B., Simojoki, A., Slade, E., Taponen, J., … & Roslin, T. (2016). Treating cattle with antibiotics affects greenhouse gas emissions, and microbiota in dung and dung beetles. Proceedings of the Royal Society B: Biological Sciences, 283(1831), 20160150. doi: 10.1098/rspb.2016.0150

43. Hanski, I., & Cambefort, Y. (1991). Competition in dung beetles. In I. Hanski & Y. Cambefort (Eds.), Dung Beetle Ecology. (pp. 305–329). Princeton, New Jersey: Princeton University Press.

44. Harry, M., Gambier, B., & Garnier-Sillam, E. (2000). Soil conservation for DNA preservation for bacterial molecular studies. European Journal of Soil Biology, 36(1), 51–55. doi; 10.1016/s1164-5563(00)00044-3

45. Hosokawa, T., Ishii, Y., Nikoh, N., Fujie, M., Satoh, N., & Fukatsu, T. (2016). Obligate bacterial mutualists evolving from environmental bacteria in natural insect populations. Nature Microbiology, 1(1), 15011. doi: 10.1038/nmicrobiol.2015.11

46. Hutchinson, G. E. (1959). Homage to Santa Rosalia or why are there so many kinds of animals?. The American Naturalist, 93(870), 145–159. doi: 10.1086/282070

47. Inoue, J. I., Oshima, K., Suda, W., Sakamoto, M., Iino, T., Noda, S., … & Ohkuma, M. (2015). Distribution and evolution of nitrogen fixation genes in the phylum Bacteroidetes. Microbes and Environments, ME14142. doi: 10.1264/jsme2.ME14142

48. Janssen, S., McDonald, D., Gonzalez, A., Navas-Molina, J. A., Jiang, L., Xu, Z. Z., … & Knight, R (2018). Phylogenetic placement of exact amplicon sequences improves associations with clinical information. MSystems, 3(3). doi: 10.1128/msystems.00021-18

49. Kikuchi, Y., Tada, A., Musolin, D. L., Hari, N., Hosokawa, T., Fujisaki, K., & Fukatsu, T. (2016). Collapse of insect gut symbiosis under simulated climate change. MBio, 7(5), e01578–16. doi: 10.1128/mbio.01578-16

50. Kingsolver, J. G., & Huey, R. B. (2008). Size, temperature, and fitness: three rules. Evolutionary Ecology Research, 10(2), 251–268.

51. Kohl, K. D., & Carey, H. V. (2016). A place for host–microbe symbiosis in the comparative physiologist’s toolbox. Journal of Experimental Biology, 219(22), 3496–3504. doi:10.1242/jeb.136325

52. Kok, R. G., Christoffels, V. M., Vosman, B., & Hellingwerf, K. J. (1993). Growth-phase-dependent expression of the lipolytic system of *Acinetobacter calcoaceticus* BD413: cloning of a gene encoding one of the esterases. Microbiology, 139(10), 2329–2342. doi: 10.1099/00221287-139-10-2329

53. Lacap, D. C., Lau, M. C., & Pointing, S. B. (2011). Biogeography of prokaryotes. In D. Fontaneto (Ed.), Biogeography of Microscopic Organisms: Is Everything Small Everywhere? (Systematics Association Special Volume Series, pp. 35–42). Cambridge: Cambridge University Press. doi:10.1017/CBO9780511974878.004

54. Legendre, P., & Anderson, M. J. (1999). Distance-based redundancy analysis: testing multispecies responses in multifactorial ecological experiments. Ecological Monographs, 69(1), 1–24. doi: 10.1890/0012-9615(1999)069[0001:dbratm]2.0.co;2

55. Lozupone, C., & Knight, R. (2005). UniFrac: a new phylogenetic method for comparing microbial communities. Applied and Environmental Microbiology, 71(12), 8228–8235. doi: 10.1128/aem.71.12.8228-8235.2005

56. Macagno, A. L., Moczek, A. P., & Pizzo, A. (2016). Rapid divergence of nesting depth and digging appendages among tunneling dung beetle populations and species. The American Naturalist, 187(5), E143–E151. doi: 10.1086/685776

57. MacArthur, R. H. (1958). Population ecology of some warblers of northeastern coniferous forests. Ecology, 39(4), 599–619. doi: 10.2307/1931600

58. Martin, M. (2011). Cutadapt removes adapter sequences from high-throughput sequencing reads. EMBnet Journal, 17(1), 10–12. doi: 10.14806/ej.17.1.200

59. Martiny, J. B. H., Bohannan, B. J., Brown, J. H., Colwell, R. K., Fuhrman, J. A., Green, J. L., … & Morin, P. J. (2006). Microbial biogeography: putting microorganisms on the map. Nature Reviews Microbiology, 4(2), 102. doi: 10.1038/nrmicro1341

60. Matsen, F. A., Hoffman, N. G., Gallagher, A., & Stamatakis, A. (2012). A format for phylogenetic placements. PLoS ONE, 7(2), e31009. doi: 10.1371/journal.pone.0031009

61. Matsen, F. A., Kodner, R. B., & Armbrust, E. V. (2010). pplacer: linear time maximum-likelihood and Bayesian phylogenetic placement of sequences onto a fixed reference tree. BMC Bioinformatics, 11(1), 538. doi: 10.1186/1471-2105-11-538

62. McMurdie, P. J., & Holmes, S. (2013). phyloseq: an R package for reproducible interactive analysis and graphics of microbiome census data. PloS ONE, 8(4), e61217. doi: 10.1371/journal.pone.0061217

63. Mirarab, S., Nguyen, N., & Warnow, T. (2012). SEPP: SATé-enabled phylogenetic placement. In Biocomputing 2012 (pp. 247–258). doi: 10.1142/9789814366496_0024

64. Moeller, A. H., Peeters, M., Ndjango, J. B., Li, Y., Hahn, B. H., & Ochman, H. (2013). Sympatric chimpanzees and gorillas harbor convergent gut microbial communities. Genome Research, 23(10), 1715–1720. doi: 10.1101/gr.154773.113

65. Moeller, A. H., Suzuki, T. A., Lin, D., Lacey, E. A., Wasser, S. K., & Nachman, M. W. (2017). Dispersal limitation promotes the diversification of the mammalian gut microbiota. Proceedings of the National Academy of Sciences, 114(52), 13768–13773. doi: 10.1073/pnas.1700122114

66. Morales-Jiménez, J., de León, A. V. P., García-Domínguez, A., Martínez-Romero, E., Zúñiga, G., & Hernández-Rodríguez, C. (2013). Nitrogen-fixing and uricolytic bacteria associated with the gut of *Dendroctonus rhizophagus* and *Dendroctonus valens* (Curculionidae: Scolytinae). Microbial Ecology, 66(1), 200–210. doi: 0.1007/s00248-013-0206-3

67. Oksanen, J., Blanchet, F.G., Friendly, M., Kindt, R., Legendre, P., McGlinn, D., … & Wagner, H. (2019). vegan: Community Ecology Package. R Package version 2.5-6. Retrieved from http://CRAN.Rproject.org/package=vegan

68. Palmer-Young, E. C., Raffel, T. R., & McFrederick, Q. S. (2018). Temperature-mediated inhibition of a bumblebee parasite by an intestinal symbiont. Proceedings of the Royal Society B, 285(1890), 20182041. doi: 10.1101/360263

69. Parker, E. S., Newton, I. L., & Moczek, A. P. (2020). (My Microbiome) Would Walk 10,000 miles: Maintenance and Turnover of Microbial Communities in Introduced Dung Beetles. Microbial Ecology, 1–12. doi: 10.1007/s00248-020-01514-9

70. Parker, E. S., Dury, G. J., & Moczek, A. P. (2019). Transgenerational developmental effects of species-specific, maternally transmitted microbiota in Onthophagus dung beetles. Ecological Entomology, 44(2), 274–282, doi: 10.1111/een.12703

71. Peres-Neto, P. R., Legendre, P., Dray, S., & Borcard, D. (2006). Variation partitioning of species data matrices: estimation and comparison of fractions. Ecology, 87(10), 2614–2625.

72. Price, D. L. & May, M. (2009). Behavioral ecology of Phanaeus dung beetles (Coleoptera: Scarabaeidae): review and new observations. Acta Zoológica Mexicana (ns), 25(1), 211–238. doi: 10.21829/azm.2009.251621

73. R Core Team (2020). R: A language and environment for statistical computing. Vienna, Austria: R Foundation for Statistical Computing. https://www.R-project.org/

74. Rosa, E., Minard, G., Lindholm, J., & Saastamoinen, M. (2019). Moderate plant water stress improves larval development, and impacts immunity and gut microbiota of a specialist herbivore. PLoS ONE, 14(2), e0204292. doi: 10.1371/journal.pone.0204292

75. Russell, J. A., & Moran, N. A. (2006). Costs and benefits of symbiont infection in aphids: variation among symbionts and across temperatures. Proceedings of the Royal Society B: Biological Sciences, 273(1586), 603–610. doi: 10.1098/rspb.2005.3348

76. Schoener, T. W. (1974). Resource partitioning in ecological communities. Science, 185(4145), 27–39. doi: 10.1126/science.185.4145.27

77. Schwab, D. B., Riggs, H. E., Newton, I. L., & Moczek, A. P. (2016). Developmental and ecological benefits of the maternally transmitted microbiota in a dung beetle. The American Naturalist, 188(6), 679–692. doi: 10.1086/688926

78. Sepulveda, J., & Moeller, A. H. (2020). The effects of temperature on animal gut microbiomes. Frontiers in Microbiology, 11. doi: 10.3389/fmicb.2020.00384

79. Shafquat, A., Joice, R., Simmons, S. L., & Huttenhower, C. (2014). Functional and phylogenetic assembly of microbial communities in the human microbiome. Trends in Microbiology, 22(5), 261–266. doi: 10.1016/j.tim.2014.01.011

80. Sheldon, K. S., & Tewksbury, J. J. (2014). The impact of seasonality in temperature on thermal tolerance and elevational range size. Ecology, 95(8), 2134–2143. doi: 10.1890/13-1703.1

81. Shukla, S. P., Sanders, J. G., Byrne, M. J., & Pierce, N. E. (2016). Gut microbiota of dung beetles correspond to dietary specializations of adults and larvae. Molecular Ecology, 25(24), 6092–6106. doi: 10.1111/mec.13901

82. Šidák, Z. (1967). Rectangular confidence regions for the means of multivariate normal distributions. Journal of the American Statistical Association, 62(318), 626–633. doi: 10.1080/01621459.1967.10482935

83. Simmons, L. W., & Ridsdill-Smith, T. J. (2011). Reproductive competition and its impact on the evolution and ecology of dung beetles. Ecology and evolution of dung beetles. Oxford: Blackwell Publishing, 1–20. doi: 10.1002/9781444342000.ch1

84. Snell-Rood, E. C., Burger, M., Hutton, Q., & Moczek, A. P. (2016). Effects of parental care on the accumulation and release of cryptic genetic variation: review of mechanisms and a case study of dung beetles. Evolutionary Ecology, 30(2), 251–265. doi: 10.1007/s10682-015-9813-4

85. Soil Science Division Staff. (2017). Examination and Description of Soil Profiles. In C. Ditzler, K. Scheffe, and H.C. Monger (Eds.), Soil Survey Manual, USDA Handbook 18 (pp. 83–233). Washington, D.C.: Government Printing Office.

86. Sun, X., Yang, Y., Zhang, N., Shen, Y., & Ni, J. (2015). Draft genome sequence of *Dysgonomonas macrotermitis* strain JCM 19375T, isolated from the gut of a termite. Genome Announcements, 3(4). doi: 10.1128/genomea.00963-15

87. Tiede, J., Scherber, C., Mutschler, J., McMahon, K. D., & Gratton, C. (2017). Gut microbiomes of mobile predators vary with landscape context and species identity. Ecology and Evolution, 7(20), 8545–8557. doi: 10.1002/ece3.3390

88. Walters, A. W., Hughes, R. C., Call, T. B., Walker, C. J., Wilcox, H., Petersen, S. C., … & Chaston, J. M. (2020). The microbiota influences the *Drosophila melanogaster* life history strategy. Molecular Ecology, 29(3), 639–653. doi: 10.1111/mec.15344

89. Wang, Y., Kapun, M., Waidele, L., Kuenzel, S., Bergland, A. O., & Staubach, F. (2020). Common structuring principles of the *Drosophila melanogaster* microbiome on a continental scale and between host and substrate. Environmental Microbiology Reports, 12(2), 220–228. doi: 10.1111/1758-2229.12826

90. Wang, Z., Xin, X., Shi, X., & Zhang, Y. (2020). A polystyrene-degrading *Acinetobacter bacterium* isolated from the larvae of *Tribolium castaneum*. Science of the Total Environment, 138564. doi: 10.1016/j.scitotenv.2020.138564

91. Whittaker, R. H. (1960). Vegetation of the Siskiyou mountains, Oregon and California. Ecological Monographs, 30(3), 279–338. doi: 10.2307/1948435

92. Wilck, N., Matus, M. G., Kearney, S. M., Olesen, S. W., Forslund, K., Bartolomaeus, H., … & Vvedenskaya, O. (2017). Salt-responsive gut commensal modulates T H 17 axis and disease. Nature, 551(7682), 585. doi: 10.1038/nature24628

93. Yilmaz, P., Parfrey, L. W., Yarza, P., Gerken, J., Pruesse, E., Quast, C., … & Glöckner, F. O. (2014). The SILVA and “all-species living tree project (LTP)” taxonomic frameworks. Nucleic Acids Research, 42(D1), D643–D648. doi: 10.1093/nar/gkt1209

94. Zhang, B., Leonard, S. P., Li, Y., & Moran, N. A. (2019). Obligate bacterial endosymbionts limit thermal tolerance of insect host species. Proceedings of the National Academy of Sciences, 116(49), 24712–24718. doi: 10.1073/pnas.1915307116

